# Archaeological Bone Assemblages Trace Changes in Turtle and Tortoise Diversity in Continental Southeast Asia

**DOI:** 10.1101/2025.03.27.645446

**Authors:** Corentin Bochaton, Wilailuck Naksri, Sirikanya Chantasri, Melada Maneechote, Supalak Mithong, Sophady Heng, Hubert Forestier, Prasit Auetrakulvit, Valéry Zeitoun, Jutinach Bowonsachoti, Julien Claude

## Abstract

Among tropical regions, continental Southeast Asia is a major hotspot of biodiversity. However, due to rapid urban expansion and severe deforestation, this area is recognized as one of the top global conservation priorities. As in other tropical regions, the prevailing idea is that the decline of local biodiversity in this region is a direct result of the overwhelming environmental changes that have occurred over the last 80 years. However, paleoecological data indicate that tropical forests have been exploited by human populations since at least the Late Pleistocene, and forest clearance for farming began several thousand years ago. To gain a complete overview of the long-term evolution of vertebrate fauna in continental Southeast Asia, it is necessary to investigate the past. This task is currently out of reach, mainly due to the lack of paleontological and historical studies regarding small animals, including emblematic taxa like turtles. Indeed, although some evidence suggests that the biodiversity of turtles in continental Southeast Asia may have changed over the last millennia, the paucity of historical records and archaeological assemblages studies currently precludes further assessment of the potential effect of past environmental changes, including potential human disturbances. The present study consists of a paleontological analysis of turtle bone assemblages collected from four prehistoric sites in Thailand and Cambodia. The aim is to test the hypothesis of changes in turtle biodiversity over the last 10,000 years. Our results indicate that turtle assemblages in most areas have experienced only limited changes across the Holocene, except for Laang Spean in Cambodia. However, a few rare taxa may represent extirpated or extinct species of Testudinidae in continental Southeast Asia at an unknown time during the Holocene. We also observed significant changes in the geographic distribution of some genera, such as *Batagur*, *Cyclemys*, and *Amyda*. The fact that turtle fauna has been rather resilient and unchanged for several millennia shows that past human impact in most investigated areas had limited effects on these taxa. The strong modifications observed in Cambodia could, however, serve as a warning about the potential fate of turtle biodiversity in this region if further loss of natural environments occurs.

## Introduction

In our modern world, the environmental degradations caused by human activities have resulted in a current rate of species extinction that is estimated to be 100 to 1,000 times greater than the “normal rate” observed throughout geological time and a genus extinction rate 35 greater (Cowie et al., 2022; Ceballos & Ehrlich, 2023). Researchers estimate that, since 1500, between 0.8% and 1.5% of vertebrate species on Earth have gone extinct (Ceballos et al., 2015; Johnson et al., 2017; Ceballos & Ehrlich, 2018). In this context, a deep understanding of the dynamics of these extinctions is critical to better evaluate the human impact on the environment. However, paradoxically, these are still not fully understood. This is particularly true for the tropics, known to be among the most threatened ecosystems. One of the main gaps in our knowledge concerns the poor understanding the scientific community has of the deep history of human impact on ecosystems and biodiversity in these regions. Notably, modern tropical rainforests are often seen as wild areas, never having been impacted by preindustrial societies and removed from human exploitation (Denevan, 1992; Willis et al., 2004; Barlow et al., 2012; Roberts, 2019). However, increasing numbers of paleoenvironmental and archaeological studies are demonstrating that this view is highly romanticized and misleading: these environments have been extensively exploited and modified by human groups since prehistory (Denham et al., 2003; Willis et al., 2004; Roberts, 2019; Stephens et al., 2019). An evaluation of the evolution of the biodiversity in these regions across the Quaternary is thus needed to clarify whether or not human populations had a major impact on it.

Among tropical areas, Southeast Asia is a major hotspot of biodiversity, hosting at least 6,220 non-fish vertebrate species (Myers et al., 2000). Due to rapid urban expansion and strong deforestation, this geographic area is recognized as one of the worldwide top conservation priorities (Sodhi et al., 2010; Duckworth et al., 2012). As a result of these anthropogenic disturbances, Southeast Asia has the highest proportion of its vascular plant, reptile, bird, and mammal species categorized as globally threatened on the IUCN Red List (Sodhi et al., 2010). Within Southeast Asia, Thailand, and Cambodia are informative areas to study past ecological phenomena in continental Southeast Asia. These countries are also among the places with the highest biodiversity and greatest human-induced threats of extinction. Both countries are located in one of the 25 identified Earth biodiversity hotspots: the Indo-Burma area, with Thailand also part of the Sundaland area (south of its peninsula) (Myers et al., 2000). Combined, these two hotspots host 3,985 non-fish vertebrate species, representing 14% of non-fish vertebrate total biodiversity on our planet. Thailand and Cambodia respectively host 1,923 and 1,028 non-fish vertebrate species, among which 9.6% and 10.4% are considered currently threatened by the IUCN (www.iucnredlist.org). However, although the animal biodiversity of Southeast Asia is currently declining and could decline even more by 2050 (Trisurat et al., 2010; Paradis, 2018), the overall environmental situation presents strong variations between the different countries, and this tendency could be reversed should the appropriate measures be taken (Pomoim et al., 2022; Botterill-James et al., 2024). As in other tropical areas, the prevailing idea is that the decline of biodiversity in Thailand is a direct effect of the modern economic revolution, which started around 1950 (Falkus, 1991), with the beginning of broad-scale agricultural exploitation (especially for rice and teak wood). These recent ecological impacts led to the loss of most of Thailand’s forest cover in less than forty years (Ramitanondh, 1989; FAO, 2020a). The situation appears similar in Cambodia, even if this phenomenon started later, around 1970 (Tsujino et al., 2019). Overwhelming environmental changes have obviously occurred in the last 80 years in continental Southeast Asia, and these changes have had a strong impact on the native vertebrate biodiversity (Baker & Phongpaichit, 2014). However, paleoecological data indicate that tropical forests have been exploited by human populations since the Late Pleistocene at least (Conrad, 2015). Likewise, the first traces of mass forest clearance for farming started around 2000 B.C. (White et al., 2004). Investigating the deeper past might improve our understanding of biodiversity loss dynamic in continental Southeast Asia.

Moreover, little is known about the past large mammal diversity of Southeast Asia (e.g., Zeitoun et al., 2010; Conrad, 2015), and almost nothing is known about the past diversity of the small animals accounting for 97% of the current terrestrial vertebrate biodiversity of this region (Chaimanee, 1998; Bochaton et al., 2024). Among these small vertebrate species are turtles and tortoises, animals that are, for half of them, considered strongly exposed to human disturbance and at risk of extinction on a global scale (Böhm et al., 2013). These animals present significant diversity in Thailand and Cambodia, with 27 and 14 species of terrestrial and freshwater turtles respectively (http://www.iucnredlist.org). However, nearly no data exist about the evolution of their diversity over the last 10,000 years, although preliminary data have begun to emerge, suggesting that part of that diversity might have been impacted by human-driven past environmental changes. Indeed, a study from the central plain of Thailand has shown that turtles and tortoises were more diverse during historical times than they are today (Claude et al., 2019). In that pioneering study, eleven taxa were identified in several archaeological assemblages dated between 4,000 and 1,000 BP. These taxa include *Batagur affinis*, *Manouria emys*, and *Geochelone* sp., three genera that are no longer present in that area nowadays. Although not figured and represented by a single individual, *Siebenrockiella crassicolis* may also have been identified in the northernmost part of Thailand in the Late Pleistocene/Early Holocene site of Spirit Cave (Conrad et al., 2016), a place where it does not seem to occur anymore. These few pieces of evidence suggest that the biodiversity of turtles in continental Southeast Asia has potentially changed over the last millennia, but the lack of occurrence data collected from archaeological assemblages currently precludes further assessment of this potentially effect of past environmental changes, including potential past human disturbances.

The present study aims to investigate this question further by conducting an in-depth paleontological study of the turtle bone assemblage collected from four prehistoric sites in Thailand and Cambodia to test the hypothesis of changes in turtle faunas over the last 10,000 years. This material was first examined in a zooarchaeological study focused on the species representing most of the assemblages, *Indotestudo elongata* (Bochaton et al., 2023). In the present work, we will focus on the investigation of the other species found in the assemblages and will discuss their identification in greater detail than is possible in a classical zooarchaeological study.

## Material and Methods

### Taxonomic *identification of the bone remains*

The archaeological assemblages of bones studied here correspond to all the non-*Indotestudo* turtle remains identified in the framework of a general zooarchaeological investigation of various bone samples (Bochaton et al., 2023). During this study, key bones presenting morphological characteristics compatible with Trionychidae, Geoemydidae, and non-*Indotestudo* Testudinidae species were photographed for further comparisons and subjected to more detailed morphological and taxonomic analysis. The identification of these bones was performed based on direct comparison with pictures and 3D models of most of the turtle species currently occurring in Southeast Asia (see specimen list in Appendix 1). This process was complicated by the fact that, as with all zooarchaeological assemblages, most of the studied material consisted of isolated bones and plates that were more or less altered by post-mortem manipulations by archaeological human populations or natural post-depositional processes. These factors made the identification of this material more challenging than that of samples corresponding to at least partially complete specimens. Consequently, precise identification at the species level was possible for only a minor proportion of the complete bone assemblage. Thus, with few exceptions, the taxa identified in the assemblages are represented by a very limited number of remains bearing potentially discriminant osteological criteria of taxonomic value. Such criteria were difficult to define because only a few studies have been conducted on the identification of isolated plates of Southeast Asian turtle species (Pritchard et al., 2009; Claude et al., 2019). We therefore had to define new identification criteria based on a sample of modern specimens gathered from several museum institutions (see specimen list in Appendix 1). Unless explicitly stated in the text, all statements regarding the morphological specificity of each taxon are made within the framework of a comparison with the species present in our reference sample only. This means that if we state a character is unique to a particular species, it is unique within the sample of modern species we studied, which includes nearly all Southeast Asian turtle species but not all turtles globally. These osteological characters were defined following the nomenclatures of Zangerl (1969) for the carapace and Szczygielski and Piechowski (2023) for the postcranial elements. We also created drawings of the carapaces and plastrons of as many Southeast Asian Testudinidae and Geoemydidae taxa as we could find in museum collections and the literature. These drawings are included in Appendix 2 and although we were not able to figure all the taxa, we hope that this initiative will facilitate future studies in the region. The reader should however keep in mind that these figures do not represent the full variability of the specimens included in our comparison dataset (see Appendix. 1).

### The studied archaeological assemblages

The studied deposits were described in detail in a previous publication (Bochaton et al., 2023). They include the Doi Pha Kan rock shelter located in northern Thailand (E 99° 46’ 37.2’’; N 18° 26’ 57.0’’), where three Hoabinhian burials dated between 11,170 ± 40 and 12,930 ± 50 BP were found (Imdirakphol et al., 2017; Zeitoun et al., 2019). This site also provided a rich archaeological assemblage corresponding to a Hoabinhian occupation older than the burials. The turtle sample studied includes 4,762 bone remains, with identified turtle taxa corresponding to *Indotestudo elongata* and rare other Testudinidae (71%), Geoemydidae (28%), and Trionychidae (0.6%).

The second site is Moh Khiew Cave, an archaeological rock shelter located in southern Thailand, in the Krabi Province (E 98° 55’ 49.27’’; N 08° 09’ 36.32’’). It was first excavated by S. Pookajorn between 1991 and 1994 (Pookajorn, 2001) and again in 2008 by P. Auetrakulvit (Auetrakulvit et al., 2012). The stratigraphy of the site comprises several archaeological layers dated from the Holocene to the Late Pleistocene, most of which correspond to Hoabinhian occupations. The material studied in the present work corresponds to that collected in 2008. The turtle sample from this site includes 5,740 bone remains, with identified turtle taxa corresponding to *Indotestudo elongata* (67%), Geoemydidae (32%), and Trionychidae (1%). These bones mainly come from two layers. The first layer provided two ^14^C dates: 7,520 ± 420 BP and 8,660 ± 480 BP (Auetrakulvit et al., 2012). The second layer provided two additional dates: 8,730 ± 480 BP and 9,270 ± 510 BP(Auetrakulvit et al., 2012).

The third site is Khao Ta Phlai Cave, located in southern Thailand, in the province of Chumphon (E 99° 5’ 49.08’’; N 10° 36’ 12.39’’). It has been excavated by the 12^th^ Regional Office of the Fine Arts Department, Nakhon Si Thammarat, since 2014. The turtle sample from this site includes 5,236 bone remains, with identified turtle taxa corresponding to *Indotestudo elongata* (63%), Geoemydidae (34%), and Trionychidae (1.9%). This bone sample is divided into two chronological assemblages corresponding to the Metal Ages and Neolithic periods. These layers have been dated based on the typology of the archaeological artifacts they have provided, such as lithic tools, ceramic shards, and metal objects. However, no direct dating is available yet, as the detailed study of the site is still ongoing.

The last site is Laang Spean Cave, a large cave of more than 1,000 m² located in northwest Cambodia, in the Battambang Province (E 102° 51’ 00.0’’; N 12° 51’ 00.0’’). The site was first excavated between 1965 and 1968 by R. Mourer and C. Mourer-Chauviré (Mourer-Chauviré et al., 1970; Mourer-Chauviré & Mourer, 1970). Following this initial exploration, the site underwent a new detailed archaeological excavation by H. Forestier between 2009 and 2019 (Forestier et al., 2015; Sophady et al., 2016). These excavations led to the discovery of an important undisturbed Hoabinhian layer dated between 4,785 ± 25 cal. BP and 11,000 ± 100 cal. BP as well as Neolithic sepulture and a Neolithic occupation date between 3,310 ± 29 BP and 3,370 ± 25 BP (Zeitoun et al., 2024). The turtle sample we studied from this site includes 8,749 bone remains, with identified turtle taxa corresponding to *Indotestudo elongata* and rare other Testudinidae (85%), Geoemydidae (13%), and Trionychidae (0.4%).

## Results

### Paleontological study

#### Geoemydidae Theobald, 1868

*Batagur* sp. Gray, 1856 – Figure. 01

**Material:** Two fused peripheral plates from Laang Spean. (**Table. 01**)

**Descriptions and comments:**

The two identified plates (**Fig. 01**) correspond to the right first and second peripheral plates of large size that are fused to the point where their separation line is no longer visible. In dorsal view, the overall aspect of the plates is smooth; they are relatively elongated, and their anterolateral margin is rounded and smooth. The sulci of the 1^st^ pleural, and of the 1^st^ and 2^nd^ marginal scutes are thin and well-visible. The first vertebral scute sulcus cannot be seen and was certainly included in the nuchal plate. In ventral view, a ventrodorsal depression is found after the scute sulcus. This element has been attributed to *Batagur* sp. because it is large, the dorsal margin of the marginal scute is situated well below the costo-peripheral suture, the fusion between plates (first and second peripheral plates and probably the first costal plate, of which a small fragment is visible on the dorsal margin of the fragment), and the two fused plates form a strongly arched anterolateral margin. The attribution to *Batagur* and not to *Orlitia* is based on the reduced extension of the marginal scutes, which are far from the costo-peripheral suture, and on the fact that the first vertebral scute does not cover the first peripheral plate and is restricted to the nuchal bone. These characters are valid for adult specimens that we did not figure in Appendix 2 because the suture between the plates completely disappeared at some point during the maturation of the specimen.

**Figure 01.**
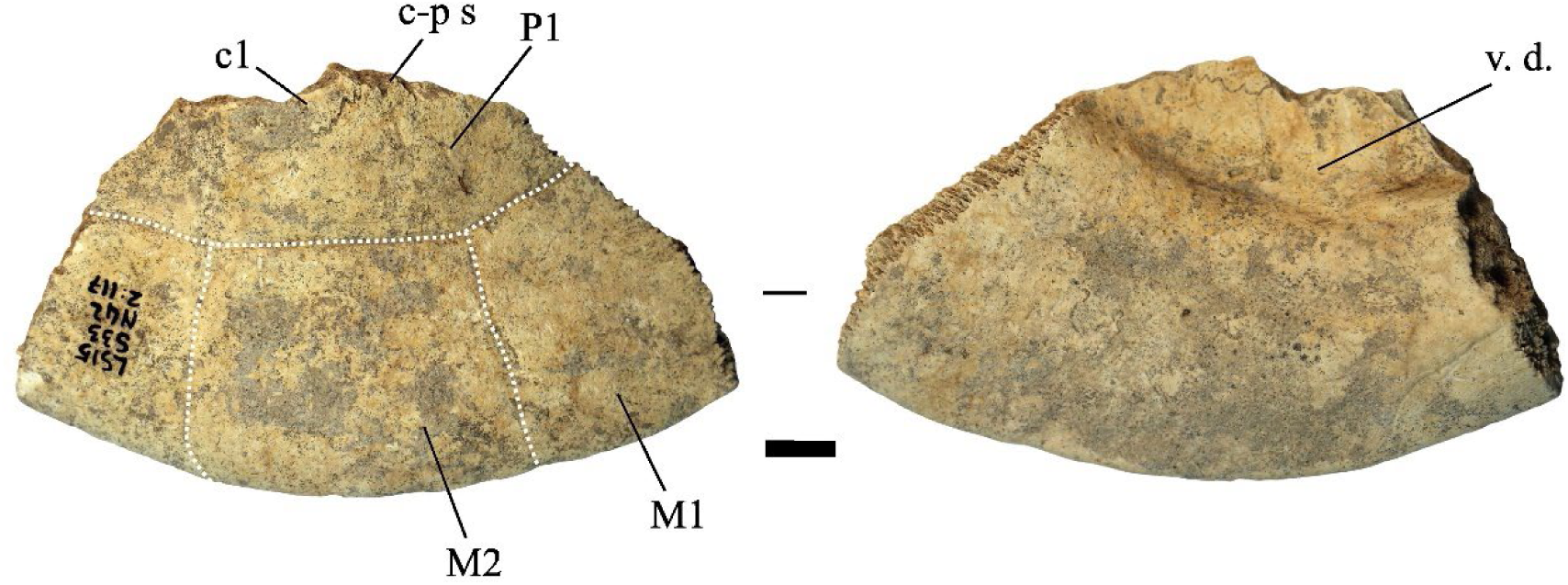
Two fused peripheral plates of *Batagur* sp. in dorsal (left) and ventral (right) views (Laang Spean). <u>Abbreviations</u>: **c1**: 1^st^ costal plate; **c-p s**: costo-peripheral suture; **v.d.**: ventrodorsal depression; **M1&2**: 1^st^ and 2^nd^ marginal scute; **P1**: 1^st^ pleural scute. Scalebars=1cm.

**Table 1.** Distribution of the identified species in the different assemblages in terms of Number of Remains. The complete account of the turtle bone remains found at the sites is published in Bochaton et al. (2023). The small discrepancy in the bone numbers corresponds to bones ultimately moved to the category of non-identified turtle bone remains between the two studies.

| Species | Doi Pha Kan | Khao Ta Phlai | Laang Spean | Moh Khiew |
| --- | --- | --- | --- | --- |
| **Geoemydidae** |  |  |  |  |
| *Batagur* sp. |  |  | 1 |  |
| *Cuora couro* | 23 | 30 | 74 | 150 |
| *Cuora* sp. |  |  |  | 1 |
| *Cyclemys* cf. *oldhamii* | 229 | 76 | 115 | 69 |
| *Heosemys annandalii* |  | 1 | 2 |  |
| *Heosemys grandis* |  |  | 5 |  |
| *Heosemys spinosa* |  | 1 |  | 16 |
| *Heosemys* cf. *grandis/annandalii* |  | 87 | 54 | 38 |
| *Malayemys* sp. | 1 | 1 |  | 3 |
| Geoemydidae indet. | 190 | 136 | 149 | 292 |
| **Testudinidae** |  |  |  |  |
| *Geochelone* sp. |  |  | 1 |  |
| *Manouria impressa* | 5 |  | 4 |  |
| *Manouria* sp. | 1 |  | 5 |  |
| Testudinidae indet. | 1 |  |  |  |
| **Trionychidae** |  |  |  |  |
| *Amyda* sp. | 3 | 5 | 2 | 7 |
| *Dogania* sp. |  |  |  | 17 |
| Trionychidae indet. | 10 | 14 | 13 | 15 |

#### *Cuora* Gray, 1856

*Cuora couro* (Leschenault de la Tour, 1812) – Figure.02 - A, D-H

**Material:** 277 remains distributed in the four sites (**Table. 01**): five nuchal plates, nine neural plates, height suprapygal plates, height pygal plates, 10 costal plates, 173 peripheral plates, 22 epiplastra, two entoplastra, five hyoplastra, 12 fragmented hyoplastra or hypoplastra, seven hypoplastra, six xiphiplastra, one scapula, and nine ilia bones.

**Description of key elements and comments:**

The plates attributed to this species present a smooth external surface with sutural contacts between the plates that are composed of many densely packed, very small pits, spikes, and crests of limited elevation and depth (**Fig. 02: B**). The plates are also thick relative to their size, and the limits between the scutes are marked by thin sulci. The significant thickness of the plates relative to their small size, the ligamentous contact between the plastron and the carapace, and the sutural contacts are characteristics we observed in *Cuora couro* and in the *C. galbinifrons* complex, and that were absent in *C. mouhotii* (**Fig. S2 and S3**).

**Figure 02.**
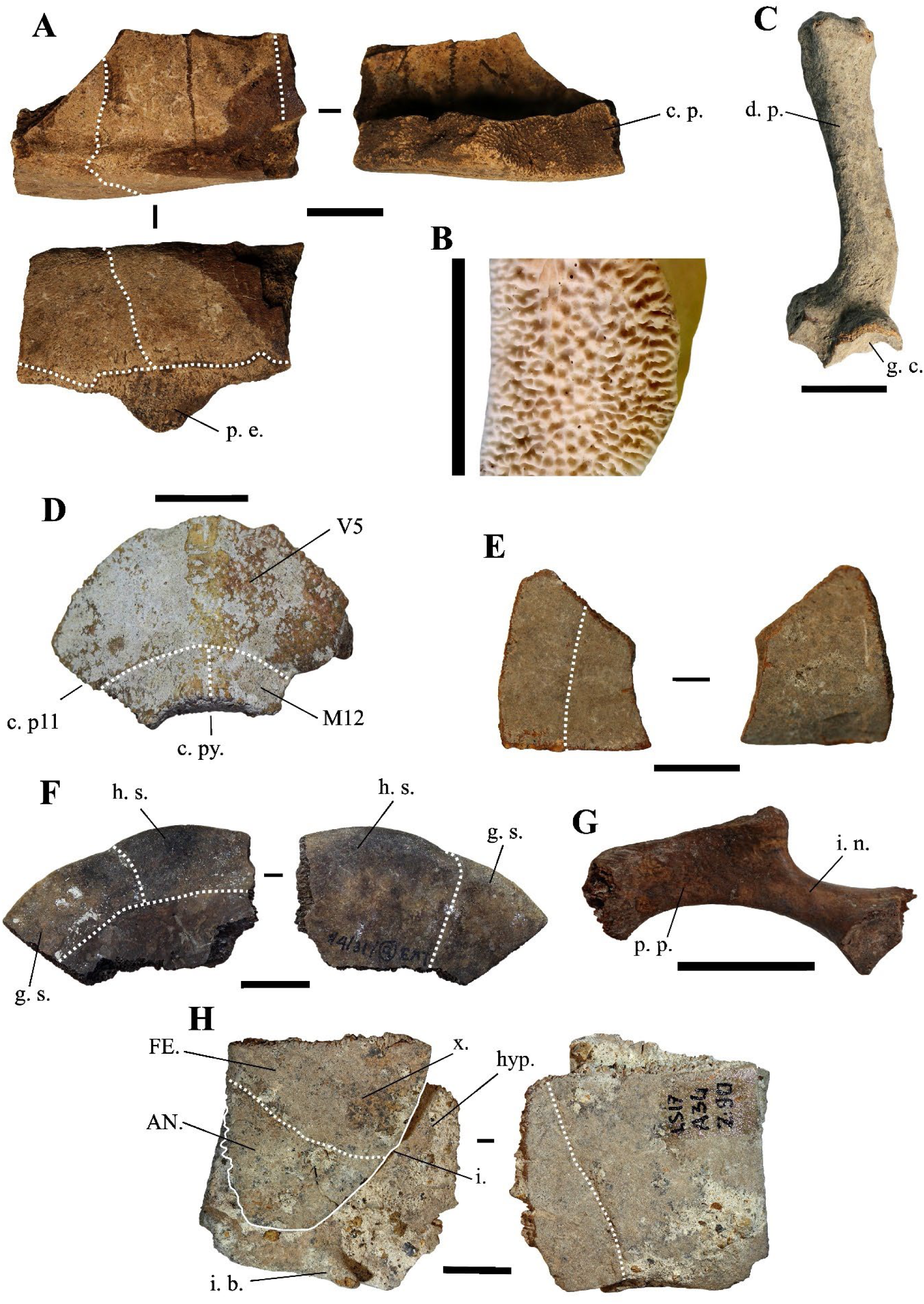
**A**) Fused 5^th^ and 6^th^ left peripheral plates of *C. couro* in lateral (top-left), medial (top-right), and ventral (bottom) views (Khao Ta Phlai); **B**) Detailed view of the contact areas between peripheral plates in a modern specimen of *C. couro*; **C**) Partial right scapula of *C. couro* in lateral view (Laang Spean); **D**) 2^nd^ suprapygal plate of *C. couro* in posterior view (Moh Khiew); **E**) Left 11^th^ peripheral plate of *C. couro* in posterior (left) and anterior (right) view (Laang Spean); **F**) Right epiplastron of *C. couro* in dorsal (left) and ventral (right) views (Moh Khiew); **G**) Right ilium bone of *C. couro* in lateral view (Doi Pha Kan); **H**) Left hypoplastron and xiphiplastron of *C. couro* cemented together by calcite (Laang Spean). <u>Abbreviations</u>: **AN.**: annal scute; **c. p.**: contact area with the plastron; **c. py.**: contact area with the pygal plate; **c. p11**: contact area with the 11^th^ peripheral plate; **d. p.**: dorsal process; **FE.**: femoral scute; **g. c.**: glenoid cavity; **g. s.**: gular scute; **h. s.**: humeral scute; **i.**: indentation; **i. b.**: inguinal buttress; **i.n.**: ilium neck; **hyp.**: hypoplastron; **M12**: 12^th^ marginal scute; **p. e.**: pointed extension; **p. p.**: posterior process; **V5**: 5^th^ vertebral scute; **x.**: xiphiplastron. Scalebars=1cm.

The nuchal plate is thick and vaulted. In dorsal view, the bone is slightly wider than long with a smooth and slightly curved anterior margin. The imprint of the cervical scute is thin and triangular with concave lateral and posterior borders. The sulcus of the anterior margin of the first pleural scute is visible on a small portion of the most lateral extremity of the plate. In ventral view, the cervical scute is broader than in dorsal view. The combination of these criteria was observed only in *C. couro* (**Fig. S2**).

The second suprapygal plate (**Fig. 02: D**) is pentagonal due to the medial contact of the two 7^th^ costal plates and sometimes hexagonal (figured specimen) in posterior view. These plates are slightly wider than high. If present, the contact with the 11^th^ peripheral plates is the same length as that of the pygal plate. The posterior sulcus of the 5^th^ vertebral scute is visible on the posterior part of the bone and is slightly convex in posterior view. The sulcus is straigth at the level of the contact with the 12^th^ marginal scutes. The second suprapygal plate also presents the above general characteristics (smooth surface, thin sulci, thickness). Within geoemydids and among our reference samples, the combination of these characteristics was found only in *C. couro* (**Fig. S3**), but it may also occur in species absent from continental Southeast Asia, namely *C. amboinensis* and *C. philippinensis*. This plate presents a morphology similar to that of *Malayemys macrocephala* and *Siebenrockiella crassicollis* (**Fig. S13, S26**), but in these species, the plates are thinner and less thick relative to their size by comparison to those of *C. amboinensis* and the *C. galbinifrons* complex (*C. galbinifrons*, *C. bourreti*, and *C. picturata*).

The peripheral plates of the bridge present a nearly smooth contact area with the plastron, indicating that the attachment of the plastron and carapace is ligamentous, with no sutural contact between the peripherals and the hypoplastron and hyoplastron. Additionally, a strong pointed extension is present on the ventromedial surface of the 5th peripheral plate (**Fig. 02: A**). This process is followed by the ligamentous contact between the hyoplastron and hypoplastron. Among our sample of modern species, these criteria have only been observed in *C. couro* and the *C. galbinifrons* complex (**Fig. S3**) but are absent in our specimen of *C. mouhotii*. This character could also be lacking in other non-Southeast Asian *Cuora* species, such as *C. aurocapitata*, or be partly absent in others, such as in *C. pani*.

The pygal plate, as the peripheral plates (**Fig. 02: E**), is small and thick. It is square in dorsal view, and its ventral margin does not exhibit any indentation. It is intersected by the medial separation of the 12^th^ marginal scutes and exhibits the general characteristics of *C. couro* (smooth surface, thin sulci, thickness, and morphology of the contact areas). In our modern sample, this morphology is only encountered in *C. couro*, while the specimens of *C. galbinifrons* we observed have a small ventral indentation (**Fig. S2**). No other Southeast Asian taxa present a thick, small-sized, square pygal plate.

The epiplastron (**Fig. 02: F**) is short, flat, and thin. Its lateral margin is smooth, regular, and rounded. In visceral view, the gular and humeral scutes cover at least half of its surface, with the area covered by each scute being nearly equal. The area occupied by these scutes presents a slightly rugged surface. In ventral view, the humeral scute covers two-thirds of the surface of the bone. The combination of these morphological criteria, along with the small size of the elements, is unique to *C. couro* and *C. galbinifrons* (**Fig. S2**). It is absent in *C. mouhotii* but might be rather general in other *Cuora* species.

The hyoplastron and hypoplastron (hypoplastron in **Fig. 02: H**) present a contact area with the carapace that is completely smooth, similar to the one described for the peripheral plates of the bridge. The inguinal and axillary buttresses are weakly developed. The contact area between the hyoplastron and hypoplastron is smooth, indicating a ligamentous connection. Among living and subfossil Geoemydidae, this morphology is, in our sample, observed only in *Cuora*. In our sample, it is unique to *C. couro* and *C. galbinifrons*, but it is absent in *C. mouhotii* (**Fig. S2, and S3**) although this might reflect a bias of our comparative sample. Indeed, this character is subject to some degree of ontogenetic variations. Ligamentous articulations between the hyo and hypoplastra have been observed in other taxa (*Cyclemys*, *Notochelys*) but the articulations are not as smooth as in *Cuora*. In addition, the femoral scute present a very short midline length, a character only observed in *C. couro*, *C. flavomarginata*, and *C. picturata* and absent in *C. galbinifrons* (Naksri et al., 2013).

The xiphiplastron (**Fig. 02: H**) is thin, flat, and wider than it is long in ventral view. In this view, its lateral margin is very convex and presents only a very small indentation at the level of the boundary between the femoral and anal scutes. It bears no trace of an anal notch, but the inter- anal sulcus is present. This morphology is unique to most species of *Cuora*, and in our sample, to *C. couro* and the *C. galbinifrons* complex (**Fig. S2**). The anal notch is also absent in *C. flavomarginata* and *C. bourreti* (Naksri et al., 2013).

The only scapula (**Fig. 02: C**) attributed to this taxon is a fragment from Laang Spean representing the dorsal process. In lateral view, this process is quite short and broad. Its orientation changes close to the glenoid cavity, as the dorsal three-fourths of its length is slightly inclined medially. Because of this, its lateral margin forms an obtuse angle. The dorsal extremity of the dorsal process is also slightly enlarged. The medial inclination of the dorsal process of the scapula is found in all the observed members of *Cuora*, but in most taxa, this process is uniformly arched, and its lateral margin forms an obtuse angle only in our specimens of *C. couro*. An enlargement of the dorsal part of the dorsal process has been seen in all the *Cuora* in our sample, but the strong build of the described fossil scapula is only observed in *C. couro*, *C. trifasciata*, and *C. galbinifrons*, while it is more elongated in the other species.

The ilium (**Fig. 02: G**) is moderately elongated. In lateral view, its ilium neck is thin and rounded, and its posterior process is enlarged and sub-quadrangular in shape. The dorsal and posterior edges of this process are rugged. This general shape of the ilium was observed in *C. galbinifrons* and *C. couro*, but in *C. galbinifrons*, the posterior process is shorter, and its posterior edge is more rounded, making the overall shape of the whole process more hook-like (see also Yasukawa, Hirayama, and Hikida, 2001). In contrast, this process is longer with more angular margins in *C. couro*.

### *Cuora* sp. – Figure. 03

**Material**: One partial nuchal plate from Moh Khiew. (**Table. 01**)

**Description and comments**:

This nuchal plate (**Fig. 03**) is only represented by its anterior portion; its posterior part is well projected in the dorsal direction. In dorsal view, the lateral sulcus of the first vertebral scute extends beyond the limits of the plate. In the same view, the cervical scute is elongated and very thin. In ventral view, the scute is triangular and approximately 2.5 times longer than wide. In our reference sample, a similar morphology of the cervical scute was only observed in *C. mouhotii* (**Fig. S3**), but in this species, although very thin, the scute is usually shorter and the anterior margin of the first vertebral scute is more convex. Additionally, the anterolateral margins of the same scute do not extend onto the adjacent plates. Outside of our references, the cervical scute morphology described here was observed as a rare morphology in several species (*Cuora couro*, *Cuora galbinifrons* complex *Cuora philippinensis*, and *C. mouhotii*) (J. C. Pers. Obs.). It thus seems likely that this plate is just an abnormal specimen of *C. couro*.

**Figure 03.**
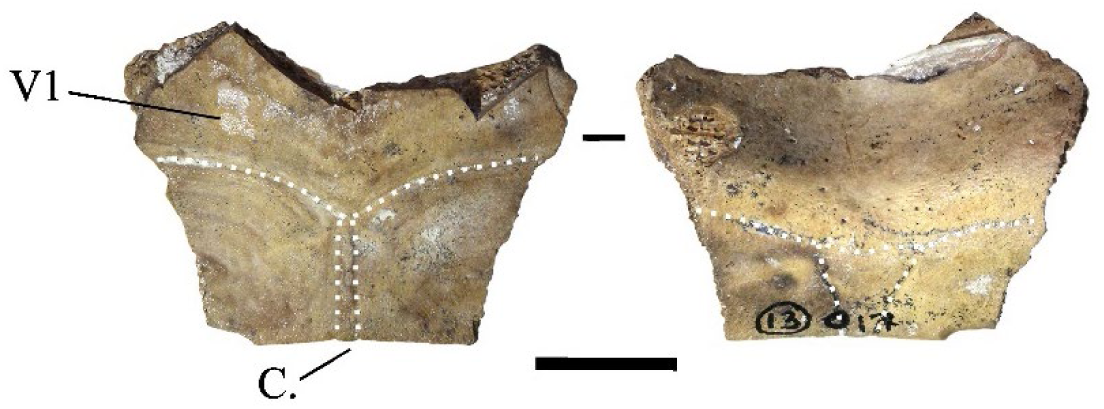
Nuchal plate of *Cuora* sp. in dorsal (left) and ventral (right) views (Moh Khiew).

#### *Cyclemys* cf. *oldhamii* Gray, 1863 – Figure. 04

**Material:** 489 remains distributed in all the sites (**Table. 01**): 20 nuchal plates, 32 costal plates, 30 neural plates, 292 peripheral plates, three 1^st^ suprapygal plates, four 2^nd^ suprapygal plates, four pygal plates, 27 epiplastra, three entoplastra, 20 hyoplastra, 18 hypoplastra, 24 fragments of hyo or hypoplastra, 12 xiphiplastra.

**Descriptions and comments:**

The nuchal plates (**Fig. 04: A, C**) exhibit a very typical morphology observed in *Cyclemys oldhamii* (**Fig. S5**). The bone is nearly as long as it is wide, with imprints of the 1^st^ pleural scutes occupying a significant part of its lateral sides in dorsal view. In the dorsal view, the cervical scute presents a slender and elongated triangular shape, laterally depressed in its anterior half and concave posteriorly. The imprint of the cervical scute is not visible in the ventral view, but this condition is variable (J. C. Pers. Obs.). In the two specimens of *Cyclemys atripons* we observed (**Fig. S4**), the cervical scute is shorter, and the bone is thickened and slightly arched below the scute, making this area very prominent. This characteristic was only observed in this species among all the Southeast Asian taxa we examined. In our six specimens of *Cyclemys dentata*, the cervical scute is shorter and a bit thicker than the morphology observed in *C. oldhamii*. We did not observe any specimens of *C. enigmatica*, *C. fusca,* and *C. pulchristriata* as these three species are currently absent from the studied area although some recent observations indicate possible occurrences of *C. enigmatica* in the Kra isthmus. Considering that some species are missing from our sample and that we had access to a limited number of comparison specimens, we propose an identification of *Cyclemys* cf. *oldhamii* while keeping in mind that the species of *Cyclemys* in the area of Moh Khiew and Khao Ta Phlai could be *C. enigmatica*.

**Figure 04.**
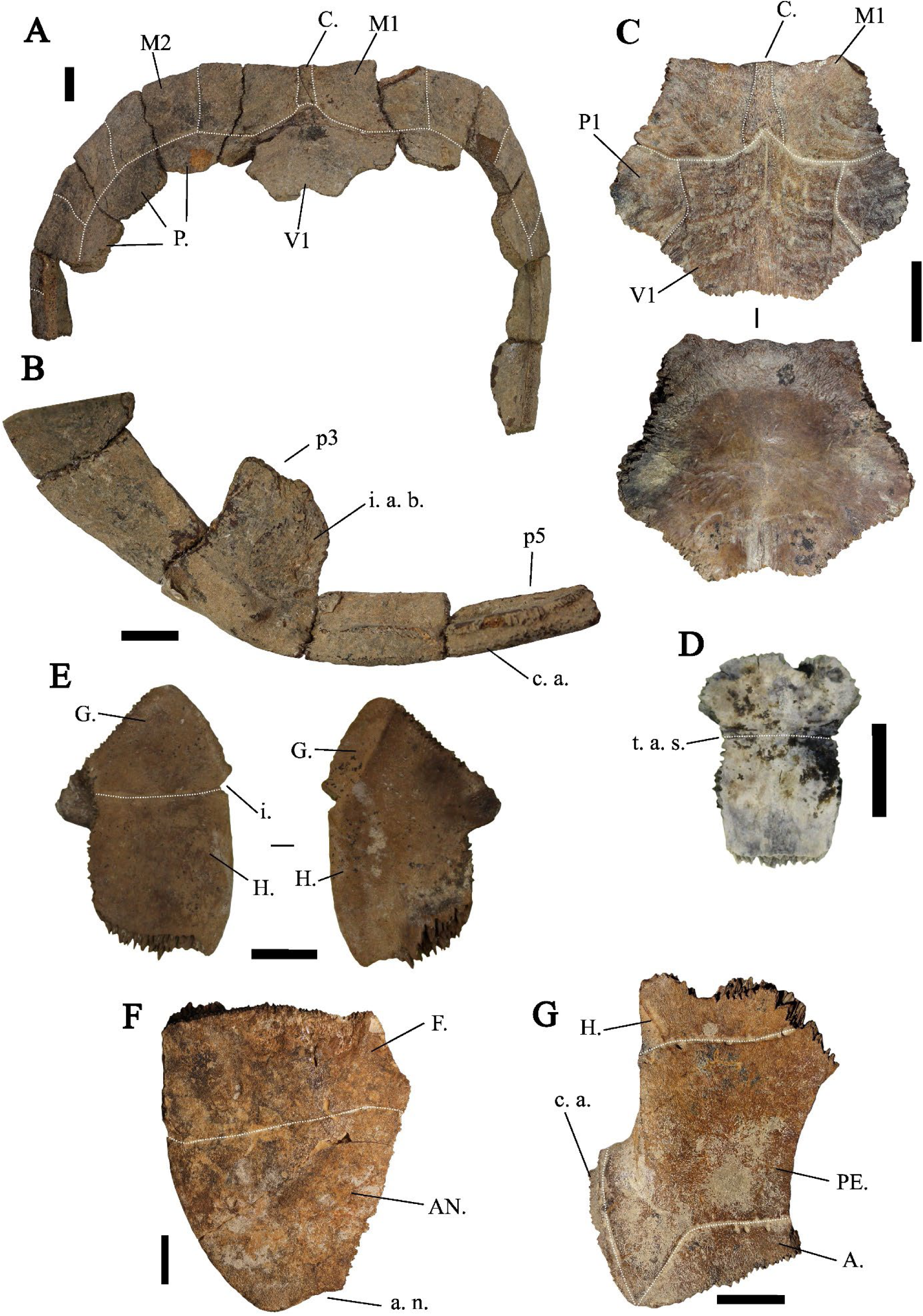
**A**) Fragments of a carapace of *Cyclemys* cf. *oldhamii* in dorsal view (Khao Ta Phlai); **B**) The plastral articular portion of the carapace illustrated in A; **C)** Nuchal plate of *Cyclemys* cf. *oldhamii* in dorsal view (Moh Khiew); **D**) 1^st^ suprapygal plate of *Cyclemys* cf. *oldhamii* (Doi Pha Kan); **E**) Epiplastron of *Cyclemys* cf. *oldhamii* in ventral (left) and dorsal (right) views (Doi Pha Kan); **F**) Xiphiplastron of *Cyclemys* cf. *oldhamii* in ventral view (Doi Pha Kan); **G)** Hyoplastron of *Cyclemys* cf. *oldhamii* in ventral view (Khao Ta Phlai). <u>Abbreviations</u>: **A.**: abdominal scute; **AN.**: annal scute; **a. n.**: annal notch; **C.**: cervical scute; **c. a.**: contact area; **F.**: femoral scute; **G.**: gular scute; **H.**: humeral scute; **i.**: indentation; **i. a. b.**: imprint of the axillary buttress; **M1**: 1^st^ marginal scute; **M2**: 2^nd^ marginal scute; **P.**: pleural scute; **PE.**: pectoral scute; **P1**: 1^st^ pleural scute; **p3**: 3^rd^ peripheral plate; **p5**: 5^th^ peripheral plate; **t. a. s.**: transverse anterior sulcus; **V1**: 1^st^ vertebral scute. Scalebars=1cm. <u>Abbreviations</u>: **C.**: cervical scute; **V1**: first vertebral scute. Scalebar=1cm.

The peripheral plates (**Fig. 04: A, B**) are thin and small in size, with the dorsal third of the height of their lateral side occupied by the pleural scutes. However, the peripheral plates of the bridge present an articulation morphology with the plastron that stands out from most other taxa. The 3^th^ to 7^th^ peripheral plates have a ventral part that projects medially with a thick contact area with the plastron, visible only in medial and ventromedial views. This contact area is marked by deep ridges mainly located in the dorsal and ventral parts of the contact area, while its center is ornamented with weakly marked ridges. The 3^rd^ to 4^th^ and 7^th^ peripherals exhibit sub-circular imprints of the ligaments attaching the carapace to the buttresses on their medial sides. The imprint of the axillary buttress can be either fully located on the third peripheral or at the intersection of the 3^rd^ and 4^th^. It can also be in contact with the lateral articular surface of the plastron or separated from it by a ridge. We have not been able to determine if this variability is of taxonomic significance at the interspecific level within the genus. The imprint of the inguinal buttress is located on the 7^th^ peripheral and might slightly overlap with the 8^th^. This attachment type is intermediate between the strong joint observed in nearly all investigated taxa, where the articular areas of the bones are strongly interlocked (as seen in the other carapace plates of *Cyclemys* species) (**Fig. S5**), and the completely smooth contact surface in some species of *Cuora* (*C. couro* and *C. galbinifrons*) (**Fig. S2**). The intermediate condition observed in *Cyclemys* only occurs in *Cuora mouhotii* (**Fig. S3**), where the attachment for the axillary buttresses is reduced and triangular.

The neural plates are thin and of trapezoid shape in dorsal view. Only a few of them, mostly from posterior positions, present traces of a keel. These bones do not exhibit characteristics unique to *Cyclemys* but have been attributed to this taxon due to its prevalence in the investigated samples and because their morphology does not differ from that of our comparative specimens of *Cyclemys oldhamii* (**Fig. S5**).

The 1^st^ suprapygal (**Fig. 04: D**) is fused with the 8^th^ neural plate. It has an elongated shape with a broad anterior part and an elongated posterior part in dorsal view. The two parts are separated by a transverse anterior sulcus between the 4^th^ and 5^th^ vertebral scutes, where the bone is slightly constricted. This morphology is common in the *Cyclemys* genus but is rare in other living geoemydid genera.

The epiplastron (**Fig. 04: E**) is thin and moderately elongated, with a circular antero-lateral margin and a well-marked acute indentation at the level of the sulcus between the humeral and the gular scutes in ventral view. The epiplastral lip is weakly developed in visceral view. Still in visceral view, the delimitation of the humeral and gular scutes is visible. This morphology is more similar to *C. oldhamii* (**Fig. S5**). Other taxa of comparable size and morphology lack a similar indentation, which can be either absent (*Cuora couro*, *Cuora mouhotii*, *Cuora trifasciata*) (**Fig. S2, and S3**) or more obtuse (*Cuora galbinifrons*, *Cyclemys atripons*, *Heosemys spinosa*) (**Fig. S4, and S9**). Some other taxa, like *Geoemyda* (**Fig. S6**), *Indotestudo*, *Melanochelys* (**Fig. S13**), *Morenia*, *Manouria* (**Fig. S11, and S12**), and *Notochelys* (**Fig. S14**), present a well-marked indentation similar to the one observed in *Cyclemys oldhamii*, but in these taxa, the anterior part of the bone forms an epiplastral projection or quadrangular shape that is absent in *C. oldhamii* (**Fig. S5**). In *Notochelys* (**Fig. S14**), the strong development of the epiplastral lips forms an indentation, absent in *Cyclemys*, at the level of the inter-gular sulcus.

The entoplastron is diamond-shaped, with its posterior part slightly more developed than the anterior one beyond the posterior humeral sulcus. Its posterior margin is subcircular in ventral view, and is crossed by the posterior sulci of the humeral scutes around its mid-length and by the posterior sulci of the gular scutes near its anterior tip. Entoplastra with similar morphologies are present in *Cuora couro* (**Fig. S2**), *Cuora galbinifrons*, and *Siebenrockiella crassicollis* (**Fig. S18**), but in these taxa, there is a more developed posterior entoplastral process in visceral view. In *Melanochelys trijuga* (**Fig. S13**) and *Geoemyda spengleri* (**Fig. S6**), the posterior sulcus of the humeral scute is closer to the posterior margin of the bone. In *Mauremys annamensis*, the entoplastron presents both of these differences compared to *Cyclemys*. In *Indotestudo elongata*, although the morphology is similar to *Cyclemys*, the entire bone is clearly thicker.

The hyoplastron (**Fig. 04: G**) is moderately elongated. In ventral view, the humero-pectoral sulcus is only visible near the anterior tip of the bone and intersects the suture line with the posterolateral part of the entoplastron. The posterior sulcus of the pectoral scute is located near the posterior margin of the bone. The articular surface with the plastron presents a morphology similar to the one described on the peripheral plates of *Cyclemys*: it is composed of a medial furrow located between two crests bearing small spikes that fit the grooves present on the peripheral plates of the bridge. This type of articular surface, also observed on the hypoplastron, is unique to *Cyclemys* among the considered modern species. In *Cuora*, although the contact between these bones is also ligamentous, the contact area in *Cyclemys* is never completely smooth. Additionally, in *Cyclemys*, the abdominal scute partly covers the hyoplastron, which is not the case in *Cuora*.

The xiphiplastron (**Fig. 04: F**) is 1.3 times longer than it is wide. Its lateral margin is smooth and circular, with a moderately developed anal notch. The femoro-anal sulcus is positioned posteriorly but close to the mid-length of the bone and runs parallel to the suture line with the hypoplastron. The combination of these characteristics is unique to *Cyclemys*. A nearly similar morphology exists in *Melanochelys trijuga* (**Fig. S13**), but the anal notch is more developed in this latter species, and the lateral lips of the anal scutes are more elevated.

#### *Heosemys* Stejneger, 1902

*Heosemys annandalii* (Boulenger, 1903) – Figure. 05 - A

**Material:** Three nuchal plates from Khao Ta Phlai and Laang Spean (**Table. 01**).

**Descriptions and comments:**

These nuchal plates (**Fig. 05: A**) are large, but their complete shape cannot be observed as only the anterior portions of the plates are preserved. The imprints of the 1^st^ pleural scutes are well visible and expanded on the postero-lateral areas of the bone. In dorsal view, the anterior margin of the bone presents a weakly marked central depression. The imprint of the cervical scute is large and heart-shaped. The attribution of these plates to *Heosemys* is based on their size, the large expansion of the 1^st^ pleural scutes on the posterolateral part of the nuchal plate, and the shape of the cervical scute. Indeed, in other large-sized taxa present in the area (*Manouria*, *Batagur*, and *Orlitia*) (**Fig. S1, S11, S12, S15, and S16**), the 1^st^ pleural scutes are either not in contact or poorly expanded on the nuchal plate, and the cervical scute is not heart-shaped. The only difference we observed between *Heosemys annandalii* (**Fig. S7**) and *H. grandis* (**Fig. S8**) in dorsal view concerns the shape of the imprint of the cervical scute on the nuchal plate. In *H. annandalii*, it is large and heart-shaped, whereas, in *H. grandis*, the scute is reduced, with a shape ranging from nearly square to a somewhat heart shape presenting a very depressed posterior margin. On their ventral surface, the nuchal plate of *H. annandalii* presents a very well-marked bulge with a strong medial depression while the same bulge in *H. grandis* is less developed and lacks the medial depression.

**Figure 05.**
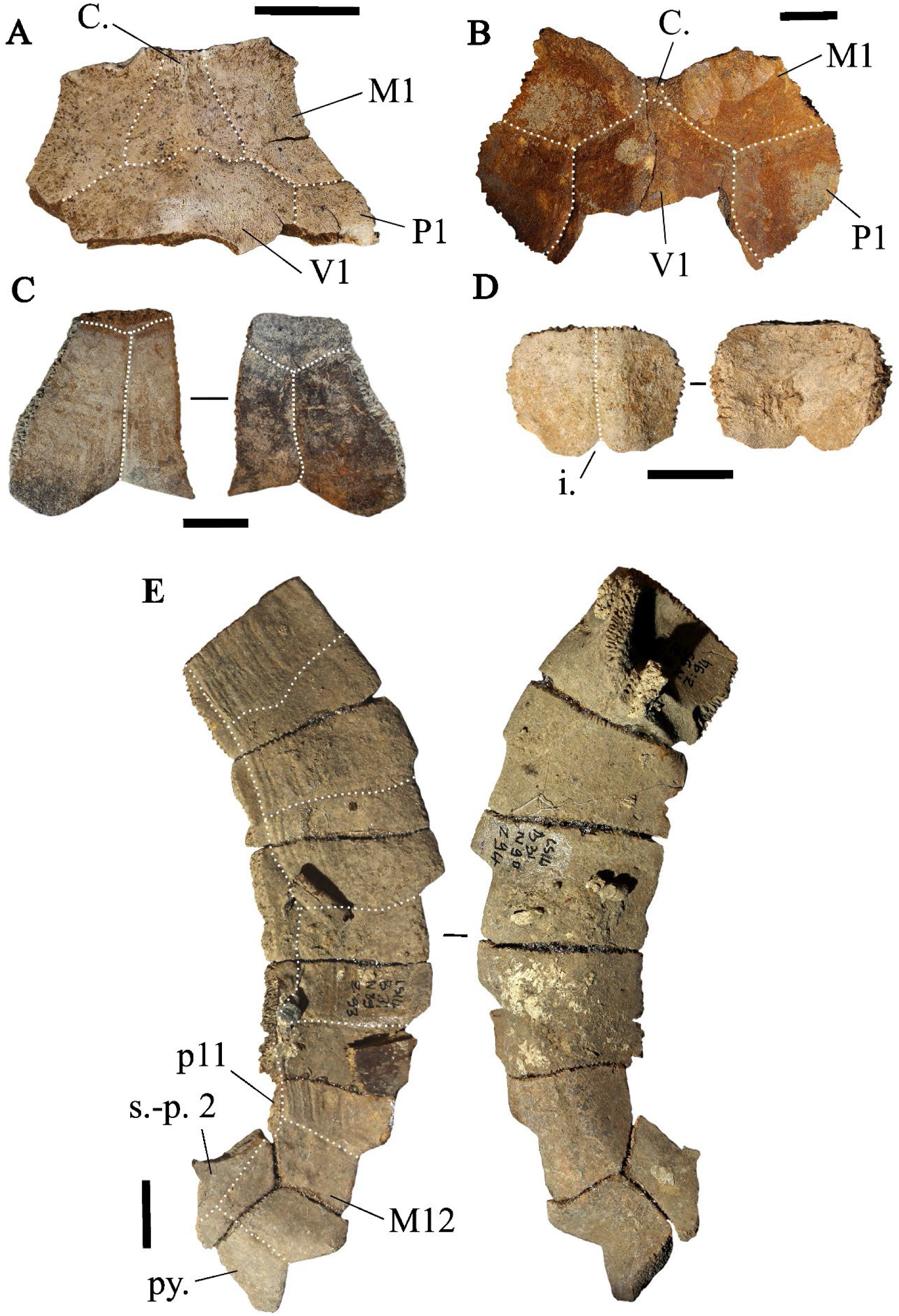
**A**) Nuchal plate of *Heosemys annandalii* in dorsal view (Khao Ta Phlai); **B**) Nuchal plate of *Heosemys grandis* in dorsal view (Laang Spean); **C**) Peripheral plate of *Heosemys grandis* in lateral view (Laang Spean); **D**) Pygal plate of *Heosemys* cf. *spinosa* in posterior view (Khao Ta Phlai); **E**) Right 7^th^ to 11^th^ peripherals, pygal and suprapygal plates attributed to *Heosemys* cf. *annandalii*/*grandis* (Laang Spean). <u>Abbreviations</u>: **C.**: cervical scute; **i.**: indentation; **M1**: 1^st^ marginal scute; **M12**: 12^th^ marginal scute; **P1**: 1^st^ pleural scute; **p11**: 11^th^ peripheral plate; **s.-p. 2**: 2^nd^ suprapygal plate; **V1**: 1^st^ vertebral scute. Scalebars=1cm.

#### *Heosemys grandis* (Gray, 1860) – Figure. 05 - B and C

**Material:** Three nuchal plates and two peripheral plates from Laang Spean (**Table. 01**).

**Descriptions and comments:**

These nuchal plates (**Fig. 05: B**) present the same characteristics as those attributed to *H. annandalii*. The only difference from the previously described morphotype concerns the shape of the cervical scute, which is reduced in the three identified specimens and nearly square in the figured one (see previous paragraph).

Posterior peripheral plates (**Fig. 05: C**) with significant serration can be attributed to this species based on their large size and morphology. Although several taxa present limited serrations on their posterior peripheral plates (9^th^ to 11^th^) (namely *Cuora mouhotii* (**Fig. S3**), *Cyclemys atripons* (**Fig. S4**), *Notochelys platynota* (**Fig. S14**), and *Heosemys annandalii* (**Fig. S7**)), strong serrations are only present in *Heosemys grandis* (**Fig. S8**), *Heosemys spinosa* (**Fig. S9**), and *Geoemyda spengleri* (**Fig. S6**). These plates were thus attributed to *H. grandis* in Laang Spean because of their large size.

#### *Heosemys* cf. *spinosa* (Gray, 1830) – Figure. 05 - D

**Material:** 15 peripheral plates and two pygal plates from Khao Ta Phlai, and Moh Khiew (**Table. 01**).

**Descriptions and comments:**

Peripheral plates have been tentatively attributed to *H. spinosa* based on their significant serration and small size (see previous comments about the peripheral plates of *H. grandis*). It is possible that these plates could correspond to young *H. grandis* specimens, but no other plates pertaining to this species were recorded at Khao Ta Phlai, and Moh Khiew.

The two pygal plates (**Fig. 05: D**) found in Khao Ta Phlai and Moh Kiew present a nearly unique morphology among the observed modern species. The plate is wider than it is high, tectiform, and has a thin, deep indentation of its ventral margin. In *H. grandis* (**Fig. S8**) and *H. annandalii* (**Fig. S7**), the posterior margin of the plate is smooth with a wide and moderately deep indentation, and the plate is not tectiform. Somehow similar morphologies were observed in the genus *Cyclemys* (**Fig. S4, and S5**), but in *H. spinosa* (**Fig. S9**) the plate is consistently wider than it is high which is not the case for the latter. Additionally, we found one of the two pygal plates in association with a series of peripheral plates from a juvenile individual that were too thick to be associated with *Cyclemys* but also lack the strong serrations observed in the other peripheral plates we attributed to *H. spinosa*. Given the limited number of elements attributed to *H. spinosa* and the potential ambiguity of the morphological characters considered in these attributions in our sample, we consider the identification of *H. spinosa* in our material as tentative.

#### Heosemys cf. grandis/annandalii – Figure. 05 - E

**Material:** 179 remains from Moh Khiew, Khao Ta Phlai, and Laang Spean (**Table. 01**): four nuchal plates, 21 neural plates, four 2^nd^ suprapygal plates, one pygal plate, four costal plates, 61 peripheral plates, five epiplastra, six entoplastra, six hyoplastra, seven hypoplastra, 30 fragmented hyoplastra or hypoplastra, nine xiphiplastra, nine humeri, six ilia bones, one ischium bone, one pubis bone, and three femora.

**Descriptions and comments:**

These bones were attributed to a large species of *Heosemys* based on their large size and the following morphological characteristics that distinguish them from the other large species considered in our comparative sample (*Batagur baska*, *Batagur borneoensis*, *Orlitia borneensis*, *Manouria impressa*, and *Manouria emys*). Regarding the general characteristics of the subfossil elements: despite their relatively large size, none of the carapace and plastron bones identified were fused to any neighboring elements. This precludes attributing these elements to *Batagur* and *Orlitia*, in which these elements tend to fuse once the animal has reached its adult size. Additionally, the subfossil elements considered here are relatively thick, which precludes an attribution of the neural and costal plates to *Manouria*, as these bones remain thin even in large individuals of this genus.

More specific observations were made on the following bones:

The 2^nd^ suprapygal plates are large, thick, hexagonal in shape, not fused to adjacent plates, and crossed by the dorsal 12^th^ marginal sulcus. The peripheral plates are thick, not fused to the adjacent plates, and high. The pleuro-marginal sulci are visible and located below the costoperipheral sutures of all the peripheral plates. A moderate serration is present exclusively along the margins of the 8^th^ to the 11^th^ peripheral plates. The epiplastron is thick, moderately elongated, and presents a moderately expanded anterior projection with a transverse anterior margin, giving the anterolateral margin of the whole bone a slightly angular shape in ventral view. Its articular surfaces with the other epiplastron and the hyoplastron are of similar lengths. The entoplastron has its ventral face crossed by the posterior gular sulcus and the posterior humeral sulcus; this characteristic is unique to *Heosemys* among the large Southeast Asian turtles. The hypoplastron is moderately elongated and presents nearly parallel anterior and posterior margins. The xiphiplastron is crossed around its mid-length by the femoro-anal sulcus, which is parallel to the straight anterior margin of the bone, and a strongly marked anal notch is present. The humerus presents the characteristics of a large Geoemydidae, but we did not identify characters specific to a given genus within this family. The shaft is moderately curved, a character shared with Trionychidae, differing from Testudinidae in which the shaft is more strongly curved. The medial and lateral processes are moderately divergent, and the deep trochanteric fossa separating them forms an angle of around 80-90°. This angle is lower in *Manouria* (around 60°) and higher in Trionychidae (around 100-110°). Also, in Trionychidae, the medial process is more individualized from the humeral head than in Geoemydidae and *Manouria*, in which the two structures are not separated. The medial process is also more extended posteriorly in Geoemydidae and Trionychidae than in *Manouria*. The ilium is antero-laterally flared (Yasukawa, Hirayama, and Hikida, 2001) as in all Geoemydidae, and its anterior part is triangular in lateral view. In *Manouria*, the anterior part of the bone is also triangular but is not antero-laterally flared. Additionally, the lateral ilial process (sensu Szczygielski and Piechowski, 2023) that is projected in a dorsolateral direction in large Geoemydidae is projected dorsally and slightly incurved in the medial direction in *Manouria*. In Trionychidae, the anterior part of the ilium is cylindrical along its entire length. Regarding the femur, in *Heosemys* and other large Geoemydidae, the trochanter major is higher than the trochanter minor and the two are coalesced, but the crest linking them is depressed in ventral view. In *Manouria*, the trochanter minor is higher than the trochanter major in ventral view, and the crest linking them is less depressed. The distal part of the bone is curved in *Manouria*, while the whole diaphysis is nearly straight in Geoemydidae. The morphology of Trionychidae is completely different, as the two trochanters are not coalesced and are separated by a deep intertrochanteric fossa, similar to the one observed on the humerus. In Trionychidae, the trochanter major is also much more developed in the dorsal direction and is triangular with a blunt apex.

#### *Malayemys* sp. Lindholm, 1931 – Figure. 06

**Material:** Three costal plates, one fragment of hyo or hypoplastron, and two fused dentaries from Khao Ta Phlai, Moh Khiew, and Doi Pha Kan. (**Table. 01**)

**Descriptions and comments:**

Three costal plates (5^th^ and 6^th^) (**Fig. 06: A**) have been attributed to *Malayemys* sp. because their medial surface presents an impression of the inguinal buttress in the ventral part of the contact area of the two plates, A faint lateral keel in the dorsal area of the plates, and a relatively straight and longitudinal vertebro-pleural sulcus. This condition is found only in *Malayemys* (**Fig. S10**) among Southeast Asian turtles. In Southeast Asia, Geoemyda has also keels, but they are located more distally, and the inguinal buttress only reaches the 5^th^ costal plate. An imprint of the inguinal buttress is also present on the 5^th^ and 6^th^ costal plates in *Batagur, Hardella,* and *Pangshura*. However, in these taxa, the buttress imprint is much stronger, covering half or more than half the length of the plates. *Geoclemys*, *Orlitia*, and *Siebenrockiella* present inguinal buttress imprints similar to *Malayemys*, but *Orlitia* and *Siebenrockiella* have mushroom-shaped vertebral scutes, resulting in non-continuous and non-longitudinal vertebro-pleural sulci. These two species also either lack (*Orlitia*) or bear lateral keels weaker than in *Malayemys* (*Siebenrockiella)*. In *Geoclemys,* the lateral keels are conversely more strongly marked than in *Malayemys*. The plates that are figured present a complex vermiculation on their lateral surface that could be either related to an antemortem phenomenon (parasitosis/senescence) or to a postmortem taphonomic alteration. Such alterations are often observed on *Malayemys* skeletal specimens.

**Figure 06.**
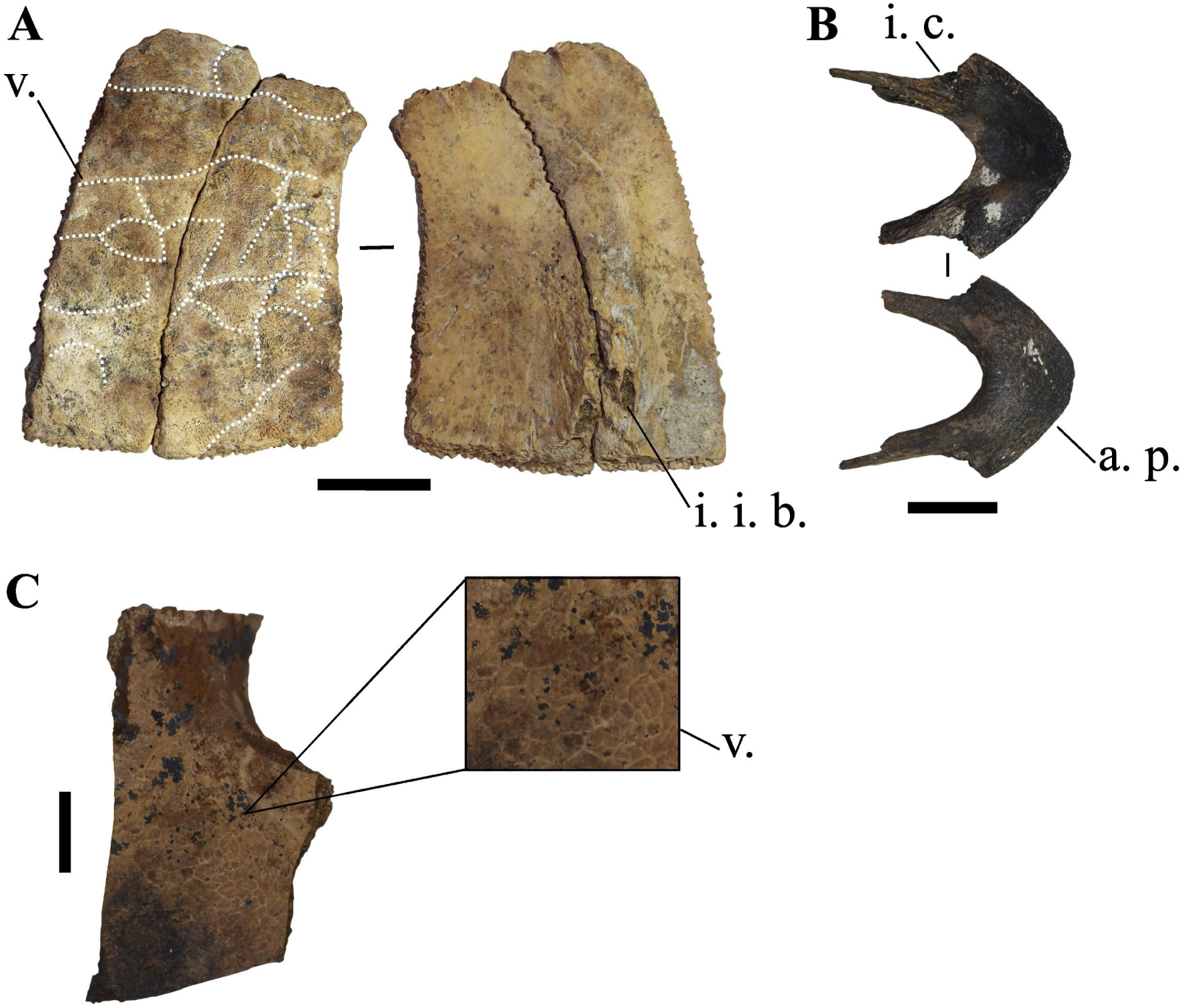
**A**) Left 5^th^ and 6^th^ costal plates of *Malayemys* sp. in dorsal (left) and ventral (right) views (Moh Khiew); **B**) Dentaries of *Malayemys* sp. in dorsal (top) and ventral (bottom) views (Moh Khiew); **C**) Hyo- or hypoplastron of *Malayemys* sp. in ventral view (Doi Pha Kan). <u>Abbreviations</u>: **a. p.**: anterior portion; **i. c.**: imprint of the coronoid bone; **i. i. b.**: impression of the inguinal buttress; **v.**: vermiculation. Scalebars=1cm.

The dentaries (**Fig. 06: B**) present no visible suture line. In dorsal view, the dentaries are short with a very wide and flat triturating surface. They present large imprints of the coronoid bones on their posterior areas, and long rami of the mandible. Long and flat lower triturating surface is known in *Malayemys*, some species of *Mauremys (M. nigricans and M. megalocephala)*, and *Geoclemys.* However, in *Geocle*mys the symphysis is shorter than in the three other species. While in *Mauremys* specimens presenting a long symphysis, the rami of the mandibles are smaller than in *Malayemys*.

A fragment of hyo- or hypoplastron (**Fig. 06: C**) has also been attributed to this genus because it presented very small anastomosed vermiculations organized in polygones on its ventral surface. Such anastomosis pattern can also be seen in *Morenia,* but in the latter, the furrows are deeper and slightly larger, while they are very thin and not deep in *Malayemys*.

#### Geoemydidae indet. Theobald, 1868

**Material:** 767 remains distributed in the four sites and representing nearly all of the skeletal parts: nine nuchal plates, 131 neural plates, 49 costal plates, 229 Peripheral plates, 11 suprapygal plates, four pygal plates, 13 epiplastra, nine entoplastra, 16 hyoplastra, 24 hypoplastra, 29 xiphiplastra, 8 undetermined fragments of plastral plates, one skull fragment, 37 scapula, 31 coracoids, 65 humeri, height radius, three ulna, 26 ilia, five ischia, 14 pubis, 38 femora, 5 tibia, 1 metapodial, and 1 phalanx. **(Table. 01)**

**Descriptions and comments:**

These bones are either fragmented or lack distinctive morphological features necessary for attribution to a specific genus. They have been classified under Geoemydidae based on their differences from elements attributed to *Indotestudo elongata* and Trionychidae. Most of these classifications are not founded on distinctive characteristics of the Geoemydidae family but are included in this study to offer a rough estimate of the family’s significance within the complete assemblage.

#### Testudinidae Batsch, 1788

*Geochelone* sp. Fitzinger, 1835– Figure. 07

**Material:** One ilium from Laang Spean. (**Table. 01**)

The ilium bone (**Fig. 07**) is partially complete and covered by a layer of calcite, with the dorsal part damaged. In lateral view, the bone is elongated, with its dorsal and ventral extremities enlarged. Its anterior and posterior edges are slightly incurved but remain nearly straight and parallel to each other in the central part of the bone. A shallow dorsal fossa is present in the dorsal area of the lateral face of the bone, though its full dorsal extent is unknown. This fossa is bordered by a thick postero-dorsal crest and a sharp antero-dorsal crest marking the anterior edge. In posterior view, the bone is slightly incurved medially, and the thick postero-dorsal crest marking the posterior delimitation of the dorsal fossa is visible. In medial view, the medial surface is smooth, and the dorsal extremity of the bone presents an angulation where the postero-dorsal crest projects laterally.

**Figure 07.**
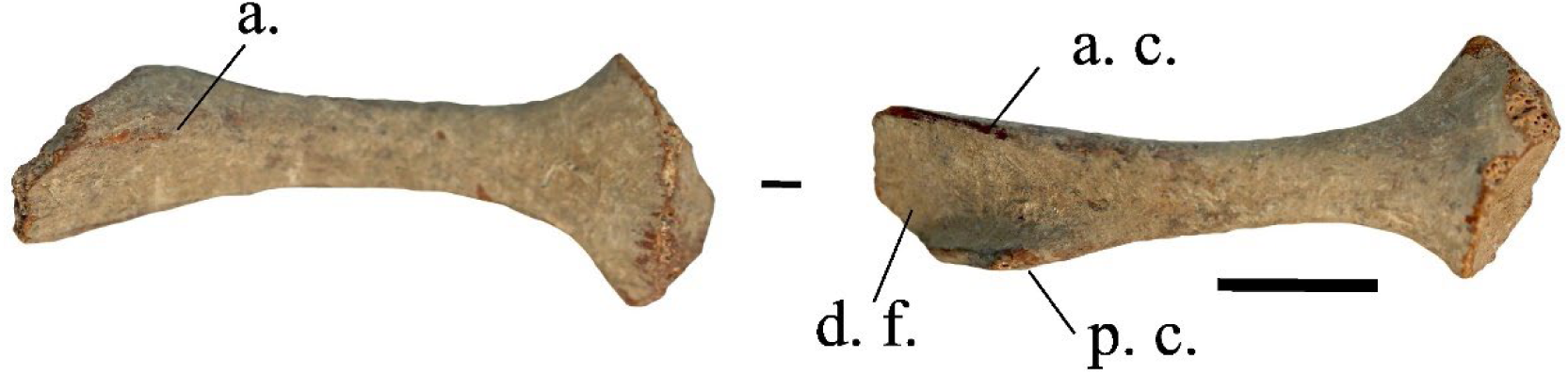
Ilium of *Geochelone* sp. in medial (right) and lateral (left) views (Laang Spean). <u>Abbreviations</u>: **a.**: angulation; **a. c.**: antero-dorsal crest; **d. f.**: dorsal fossa; **p. c.**: postero-dorsal crest. Scalebar=1cm.

The fact that the ilium presents enlargement of its dorsal and ventral extremities, a nearly straight anterior edge, and a distal fossa are characteristics of testudinoid turtles (Zug, 1971). Among all the Southeast Asian Geoemydidae we observed, the dorsal portion of the ilium presents a broad range of morphologies (also figured in Yasukawa, Hirayama, and Hikida, 2001), but its anterior and posterior edges are never close to being parallel. Among the specimens we saw, such morphology is only encountered in *Geochelone elegans* (we did not have access to a *G. platynota* specimen) and *I. elongata*. However, in the latter species, the dorsal fossa is less deep, and its limits are well-defined on the lateral surface. Additionally, *I. elongata* does not exhibit dorsal enlargement, and this seems to be the case in our fossil, even if the morphology of its dorsal part cannot be fully assessed. We thus attribute our ilium to an undefined member of the genus *Geochelone*, as we were not able to assess the inter-specific variability of the ilium within this genus. There is a possibility that this remain could be associated with the Testudinidae indet. nuchal plate described below, but since the remains were not found at the same site, this cannot be demonstrated so far.

#### Manouria impressa (Günther, 1882) – Figure. 08 - A, B, C

**Material**: Three neural plates, five costal plates, and one pygal plate from Doi Pha Kan and Laang Spean. (**Table. 01**)

**Description and comments:**

The neural plates and costal plates (**Fig. 08: A and B**) attributed to *Manouria* exhibit a distinct morphology with prominent elevations along the sulcus. In the costal plates, the anterolateral, posterolateral, and medial parts of the costal bone, separated by well-defined elevations along the sulcus, are strongly depressed. Additionally, the morphology of these plates retains a juvenile appearance as they do not contact each other along the full length of their lateral margins. The ribs are also visible on half of the length of the costal plates that do not bear the transverse sulcus of the pleural scutes. This morphology has only been observed in *Manouria impressa* (**Fig. S12**).

**Figure 08.**
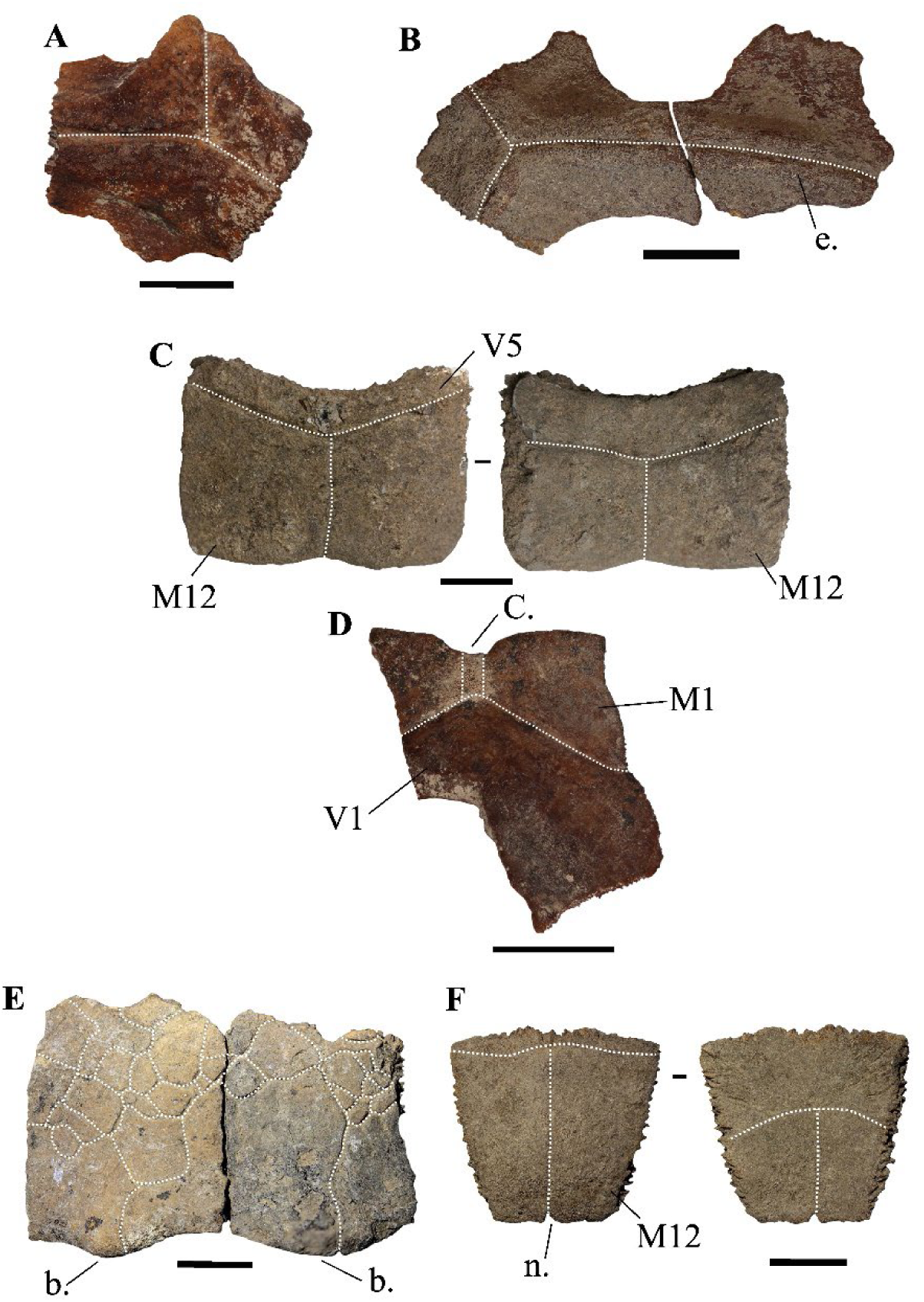
**A**) Costal plate of *Manouria impressa* in dorsal view (Laang Spean); **B**) Costal plate of *Manouria impressa* in lateral view (Doi Pha Kan); **C**) Pygal plate of *Manouria impressa* in anterior (right) and posterior view (left) (Doi Pha Kan); **D**) Nuchal plate of *Manouria* sp. in dorsal view (Doi Pha Kan); **E**) Peripheral plates of *Manouria* sp. in lateral view (Laang Spean); **F**) Pygal plate of *Manouria* sp. in anterior (right) and posterior (left) views (Laang Spean). <u>Abbreviations</u>: **b.**: bulge; **C.**: cervicale scute; **e.**: elevation; **M1**: 1^st^ marginal scute; **M12**: 12^th^ marginal scute; **n.**: notch; **V1**: 1^st^ vertebral scute; **V5**: 5^th^ vertebral scute. Scalebar=1cm.

The pygal plate (**Fig. 08: C**) is wider than it is high, with a rectangular shape in posterior view and a slightly depressed dorsal margin. It is crossed by the posterior sulcus of the 5^th^ vertebral scute, which does not overlap with the dorsal margin of the bone. Additionally, the sulcus between the 12^th^ marginal scutes is present. This morphology is specific to *M. impressa* (**Fig. S12**); on the pygal plate of *Manouria emys* (**Fig. S11**), the posterior sulcus of the 5^th^ vertebral scute overlaps with the dorsal margin of the bone.

#### *Manouria* sp. Gray, 1854 – Figure. 08 – D, E, and F

**Material:** One nuchal plate, two peripheral plates, one pygal plate, one entoplastron, and one humerus from Doi Pha Kan and Laang Spean. (**Table. 01**)

**Description and comments:**

The nuchal plate (**Fig. 08: D**) is partly broken but was likely longer than wide in dorsal view. In lateral view, the dorsal margin of the bone forms a 110° angle. In dorsal view, the plate presents long anterolateral margins crossed at their mid-length by the anterior sulcus of the 1^st^ vertebral scute. The lateral sulcus of the 1^st^ vertebral scute is not visible. The cervical scute is quadrangular and marked by a well-defined indentation of the anterior margin of the bone. In ventral view, the cervical scute is nearly square. This nuchal plate has been attributed to a juvenile individual of *Manouria* based on the combination of the following characteristics only encountered in this genus: nuchal plate longer than wide, and occurrence of a small quadrangular cervical scute (**Fig. S11, and S12**).

The two peripheral plates attributed to *Manouria* sp. (**Fig. 08: E**) present a very unusual morphology. They both have a sinusoidal ventral margin with a medial bulge. Additionally, they bear a peculiar pathological condition; their dorsal surface exhibits a scaly appearance formed of deep irregular furrows that mask the sulci of the marginal scutes. These two pieces have been associated with *Manouria* based on the observation of similar elements on a modern specimen of *Manouria* sp. However, this modern specimen did not present the pathological condition observed in these fossils.

The pygal plate (**Fig. 08: F**) is relatively thin and has a trapezoidal shape in posterior view. The posterior sulcus of the 5^th^ vertebral overlaps with the dorsal limit of the plate, the 12^th^ marginal intersulcus is present, and a small notch is visible on the central part of the ventral margin of the bone. In our comparative sample, the closest correspondence we found is *Manouria emys*. However, the pygal plate of *M. emys* is always wider than high in juvenile and adult specimens, and the shape of the bone is less triangular than our fossil. As we lack appropriate references for *M. impressa* to rule out the possibility that this plate might be attributed to a modern species of *Manouria*, we tentatively refer this plate to an undefined species of *Manouria*.

The ventral face of the only subfossil entoplastron attributed to *Manouria* sp. is crossed by the medial sulcus of the humeral and pectoral scutes but does not exhibit any other sulci. This condition has only been observed in *Manouria* (**Fig. S11, and S12**). This plate tends to be as wide as long in adult individuals but is more elongated in juvenile individuals, a condition observed in our subfossil element.

A humeral diaphysis can be associated with *Manouria* due to its large size and its morphological similarity to this taxon. It exhibits limited torsion and a well-developed medial process (*sensu* Hermanson et al., 2024) in a vertical position parallel to the main proximo-distal axis of the diaphysis.

#### Testudinidae indet. Batsch, 1788– Figure. 09

**Material:** One nuchal plate from Doi Pha Kan. (**Table. 01**)

**Descriptions and comments:**

The nuchal plate (**Fig. 09**) is pentagonal in dorsal view. In lateral view, the ventral margin of the bone forms a 110° angle as its posterior part projects dorsally. The cervical scute is absent in both dorsal and ventral views. In dorsal view, the anteromedial margins of the 1^st^ pleural scutes are visible in the posterior half of the bone. The anterior margin of the plate is notched at the level of the sulcus between the 1^st^ marginal scutes. On the unbroken right side of the plate, the 1^st^ marginal scute extends well posteriorly and nearly reaches the suture between the 1^st^ peripheral and 1^st^ costal plates. In ventral view, sulcus between the 1^st^ marginal scutes extends over more than half the length of the plate. The contact with the first peripheral plate is well-extended, and longer than the contact with the costal plate.

**Figure 09.**
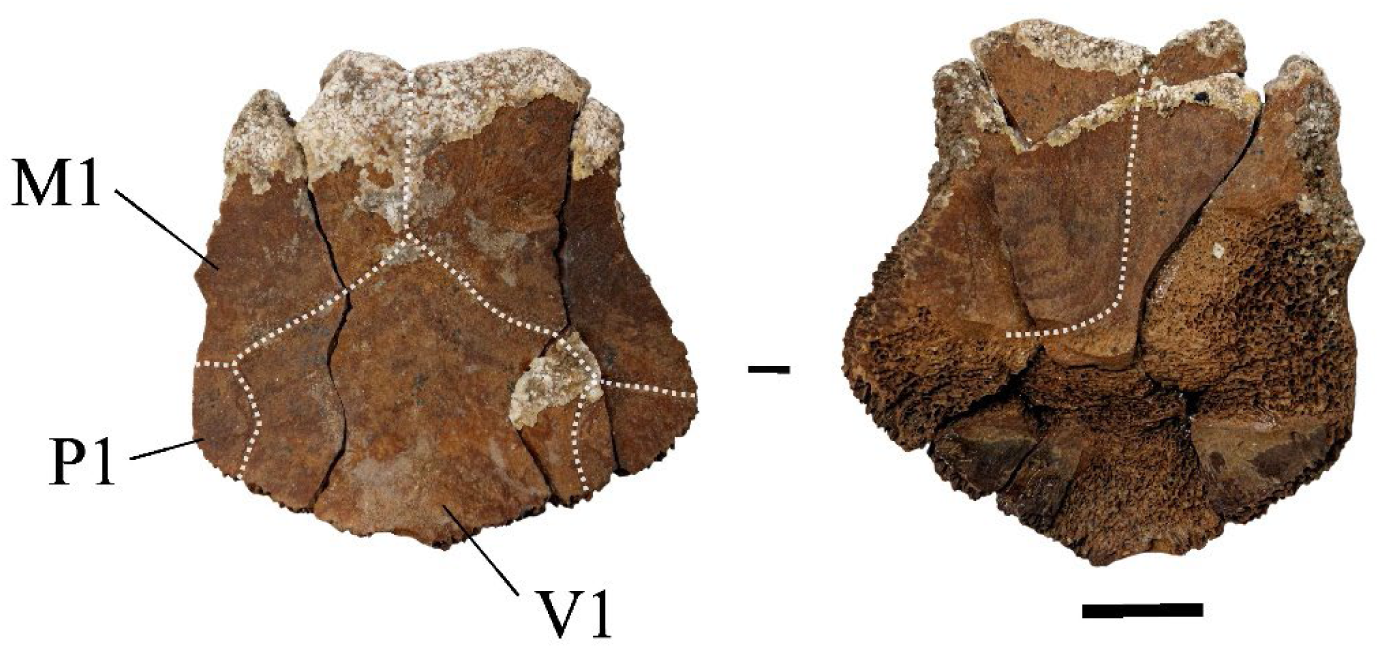
Nuchal plate of *Testudinidae* indet. in dorsal (left) and ventral (rigth) views (Doi Pha Kan). <u>Abbreviations</u>: **M1**: 1^st^ marginal scute; **P1**: 1^st^ pleural scute; **V1**: 1^st^ vertebral scute. Scalebar=1cm.

Several characters support the attribution of this fossil to the family Testudinidae: the absence of a cervical scute, the dorsal posterior extension of the marginal that nearly reaches the suture between the 1^st^ peripheral and 1^st^ costal plates, and the visceral extension of the marginal scute. The first character is found in several species of Testudinidae, while the second and third are present in all Testudinidae. On the other hand, in most Testudinidae and all its Asian forms, the pleural scutes rarely cover the nuchal to a large extent, which is a character more frequently observed in Geoemydidae. However, the absence of a cervical scute is known to occur very rarely and only in a few taxa of Geoemydidae (*Heosemys*, *Cuora*) (J. C. pers. obs.). The absence of a cervical scute is also found in Podocnemididae Cope 1869, a family currently present only in Madagascar and South America and in the tertiary fossil record of Eurasia (Rhodin et al., 2021). Some species of this group may have been present until the Late Miocene in continental Southeast Asia (Hirayama et al., 2023), but the nuchal plate is usually flat in visceral view for this group while it shows a concavity for Cryptodira. Assuming this plate belongs to a Testudinidae, the comparison with the two modern Southeast Asian species lacking a cervical scute, namely *Geochelone platynota* and *Indotestudo travancorica*, is rather inconclusive. In *I. travancorica*, the nuchal plate is flatter, its anterior margin is not strongly indented, and the posterior margin presents a well-marked concavity for the 1^st^ neural plate. In *G. platynota*, the medial contact between the 1^st^ marginals is much shorter than in our fossil, and the pleural scute does not or not strongly overlap the nuchal plate. Therefore, we attribute our nuchal plate to an unknown Testudinidae, pending for new material in the Doi Pha Kan locality.

Regarding previously published Quaternary fossils, Claude et al. (2019) mentioned the presence of another testudinid in the central plain of Thailand, different from *Manouria* and *Indotestudo*, tentatively assigning this material to *Geochelone platynota* based on geographic arguments. It is possible that the material we attribute to *Geochelone* sp. and Testudinidae indet. could be related to the same taxon. This or these extinct Testudinidae might have been present in Thailand and Cambodia during the Holocene and possibly the terminal Pleistocene.

#### Trionychidae Gray, 1825

*Amyda* sp. Geoffroy Saint-Hilaire, 1809 – Figure. 10

**Figure 10.**
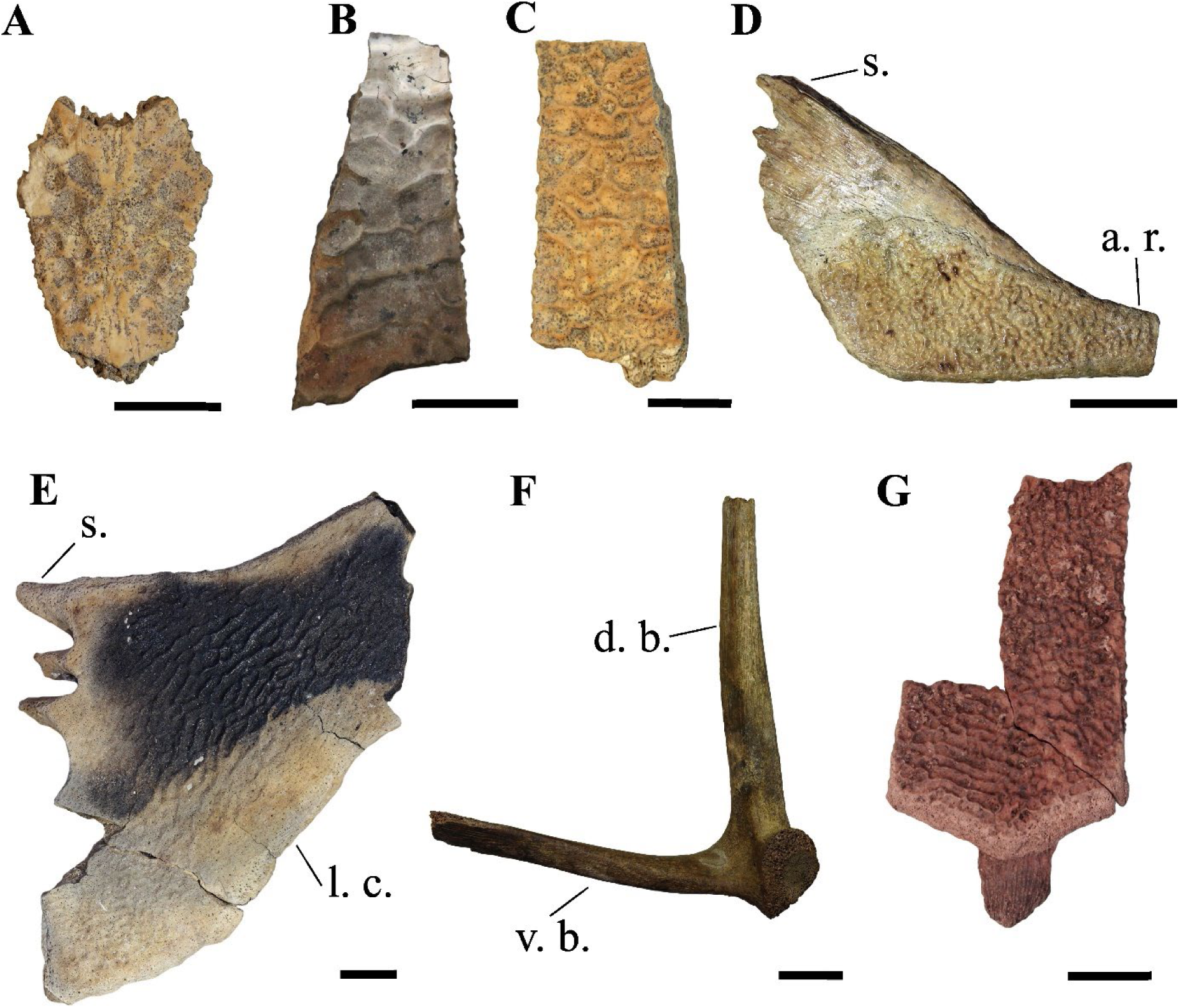
**A**) Neural plate of *Amyda* sp. in dorsal view (Laang Spean); **B**) Coastal plate of *Amyda* sp. in dorsal view (Doi Pha Kan); **C**) Coastal plate of *Amyda* sp. in lateral view (Laang Spean); **D**) hyoplastron of *Amyda* sp. in ventral view (Moh Khiew); **E**) Xiphiplastron of *Amyda* sp. in ventral view (Moh Khiew); **F**) Scapula of *Amyda* sp. in anterior view (Moh Khiew); **G**) Coastal plate of *Dogania* sp. In lateral view (Moh Khiew). <u>Abbreviations</u>: **a. r.**: axillary region; **d. b.**: dorsal branch; **s.**: spikes; **l. c.**: lateral convexity; **v. b.**: ventral branch. Scalebars=1cm.

**Material:** 15 remains from Doi Pha Kan, Khao Ta Phlai, Laang Spean, and Moh Khiew: one neural plate, 11 costal or unidentified carapace plates, one hyoplastron, one xiphiplastron, and one scapula. (**Table. 01**)

**Descriptions and comments:**

One neural plate (**Fig. 10: A**) has been tentatively attributed to *Amyda* sp. The plate is complete, measuring 2.5 cm in length, and its dorsal surface is covered with irregular pits. The maximum number of pits across the width of the plate is six, which is consistent with what is observed in other large trionychids. The pits appear shallower than those in the genus *Chitra* and do not become more elongated near the anterior or posterior margins of the bone.

Six fragments of costal or unidentified plates (**Fig. 10: B and C**) have been attributed to *Amyda* based on the morphology of their dermal ornamentation. This ornamentation consists of vermiculated sculptures and sharp ridges, both forming well-defined, irregular polygonal depressions across all external surfaces of the bones. Some elements also exhibit longitudinal ridges of higher terrain (sensu Pritchard et al. 2009). The ornamentation pattern remains consistent across all surfaces, showing little variation near the edges of the bones, except at the lateral margins of the costal plates, where the polygons become less defined and shallower. The presence of longitudinal ridges is characteristic of *Amyda* among the genera currently living in Thailand but is also observed in the southeast asian genera *Palea* and *Pelodiscus*. In contrast, the ornamentation patterns of other Thai taxa differ: in *Pelochelys* and *Chitra*, the ornamentation is less random and changes near the anterior and posterior sutures of the costal plates. Additionally, in *Pelochelys*, the ornamentation pattern remains consistent near the lateral margins of the costal plates (Pritchard et al., 2009).

A fragment of the medial side of a hyoplastron (**Fig. 10: D**) can be attributed to *Amyda* sp. The fragment is 4 cm wide, but the total width is estimated to be approximately 9 to 10 cm. The bone is narrow in its axillary region. The ventral surface is covered by a callosity ornamented with small ridges and punctuations. The medial area features three or four well-developed spikes that are closely packed and project in the same anteromedial direction. Among the taxa currently present in Thailand, this fragment can only be referred to *Amyda*. In *Pelochelys*, the axillary region of the hyoplastron is broad, while the medial spikes are either absent or poorly developed in *Dogania* and oriented in varying directions, from anteromedial to medial, in *Chitra*.

One left xiphiplastron (**Fig. 10: E**), missing a small portion of its anterior side, has been attributed to *Amyda* sp. The bone is triangular in shape, approximately 8 cm in length, with a strongly concave lateral side. On the medial side, spikes and sockets are present for articulation with the right xiphiplastron. The ventral surface features a callosity that covers most of the area and is ornamented with small, vermiculated ridges running parallel to the lateral side. This ornamentation becomes less prominent and fades near the bone’s margins. The triangular shape of the bone, the presence of medial spikes and sockets, the lateral concavity, and the morphology of the ornamentation are diagnostic of the genus *Amyda*.

One scapula (**Fig. 10: F**), with an angle of approximately 60° between its ventral and dorsal branches, has been tentatively attributed to *Amyda* due to the absence of distinguishing features separating it from this species. However, a detailed comparison with other Trionychidae taxa currently living in Thailand was not conducted.

#### *Dogania* sp. Gray, 1844 – Figure. 10

**Material:** 17 remains from Moh Khiew: one neural plate, 13 coastal plates, one hyoplastron, and two unidentified carapace plates. (**Table. 01**)

All the remains attributed to *Dogania* exhibit a very distinctive dermal ornamentation pattern (**Fig. 10: G**), which is much finer than that of *Amyda* and other Southeast Asian taxa. This pattern consists of very small, deep depressions of irregular shape, along with small tubercles located in the distal area of the costal plates (Pritchard et al., 2009).

#### Trionychidae indet. Gray, 1825

**Material:** 57 remains distributed in the four sites: 33 costal plates, one epiplastron, two unidentified carapace plates, one dentary, one coracoid, three scapula, six humeri, one radius, two ilia, one pubis, four femora, and two tibias. **(Table. 01)**

**Descriptions and comments:**

These bones correspond to either fragmented elements or anatomical parts whose morphology was not distinctive enough to be attributed to a specific genus, or for which appropriate comparative references were not available. However, they were attributed to Trionychidae based on direct comparisons with several members of this family (postcranial and cranial elements) and on the presence of ornamentation that, although not well-preserved, was insufficient to permit a subfamily-level attribution (carapace elements).

#### Distribution of the species in the investigated assemblages (Table. 01)

Regarding the Geoemydidae: the genus *Batagur* has been identified only at the site of Laang Spean in a Neolithic context. The genera *Cuora* (*Cuora couro*) and *Cyclemys* have been identified in all four assemblages. *Heosemys* have been identified in all sites except in northern Thailand (Doi Pha Kan), with two or three species represented (*H. spinosa*, *H. grandis*, and H*. annandalii*). The genus *Malayemys* was identified in Moh Khiew, Doi Pha Kan and Khao Ta Phlai. For the family Testudinidae, *Indotestudo elongata* was present in all the assemblages (Bochaton et al., 2023). Additional Testudinidae species were identified in the present study in Doi Pha Kan and Laang Spean, with *Manouria impressa*, *Manouria* sp., and at least one undetermined species attributed to Testudinidae sp. in Doi Pha Kan and to *Geochelone* sp. in Laang Spean. Regarding Laang Spean we observed no chronological tendency in the distribution of the identified species. For the Trionychidae: the genus *Amyda* was identified in Laang Spean, Khao Ta Phlai, Doi Pha Kan and Moh Khiew, and the genus *Dogania* only in Moh Khiew. Unidentified Trionychidae were also identified in the four sites.

## Discussion

### Meaning of the archaeological accumulation in terms of past biodiversity

Before discussing the significance of the data provided by paleontological analyses in terms of past biodiversity, it is necessary to address the question of the significance of these data and the mode of accumulation of the studied archaeological bone assemblages. Bone samples do not represent a frozen image of the surrounding biodiversity; they provide distorted images that reflect more on how the bone assemblage was created than the biodiversity of the surrounding animals at the site. To understand and interpret the biodiversity data provided by a bone sample, it is first necessary to understand how it was formed, in an attempt to identify, if not correct, its biases. To achieve this, taphonomic analyses and, in the case of archaeological sites, zooarchaeological analyses are needed. Such analyses were not conducted in the present study but have been performed in previous studies (Auetrakulvit, 2004; Forestier et al., 2015; Frère et al., 2018; Bochaton et al., 2019, 2023). Regarding the turtle remains, these studies unambiguously demonstrate that they correspond to animals gathered by prehistoric hunters for consumption and that they were therefore transported over some distance. The potential contribution of additional accumulation agents (i.e., birds or carnivores) is insignificant. This has major implications, as all the specimens present at the site are of relatively large size, a compositional bias that could be amplified by the bone recovery protocol in the field (see Bochaton et al., 2023). In the four sites, the hunters focused their gathering efforts on a single species, *Indotestudo elongata*. The methods that might have been used to trap these tortoises could include trapping (Santos et al., 2020; Bochaton et al., 2023), similar to the methods possibly used for other small animals like monitor lizards (Bochaton et al., 2019). The same traps could have also caught *Cuora* individuals, which are often active outside of water or can be found in shallow water. However, this cannot account for the more aquatic species, including *Cyclemys*, which is well-represented in the assemblages. This probably indicates the use of fishing techniques, corroborated by the presence of strictly aquatic fauna (mollusks and fish) in all the sites (Auetrakulvit, 2004; Forestier et al., 2015; Frère et al., 2018; Bochaton et al., 2019, 2023). Nevertheless, the exploitation of freshwater fauna was clearly secondary in all the assemblages. This could be related to the distance to freshwater sources. In addition to a likely lower representation of aquatic species, the studied bone samples clearly suffer from a near absence of large individuals. As far as turtles are concerned, this phenomenon is probably explained by the weight of these animals relative to their meat yield. Indeed, transporting these animals was probably not easy, but they are also simpler to keep captive and alive than mammals or birds and could thus be kept in the settlement for future consumption. Large animals that were impossible to move might have been excluded from the archaeological assemblages because their carcasses could have been left at the killing site, with only their meat carried to the settlement. Conversely, small animals might not have been collected because they did not represent an abundant enough source of meat to justify their transport and culinary preparation. Of course, hunting strategies could also involve less straightforward and practical considerations, such as ritual hunting (Balée, 1985), which are impossible to address with the archaeological record. Overall, it is clear that the archaeological assemblages are not a faithful representation of past turtle biodiversity, as aquatic animals are poorly represented and large and small animals are excluded. Despite these biases, it is interesting to observe that the coverage of freshwater species in the different sites seems fairly good. Indeed, the species in the sites reveal species associations similar to those observed today in most freshwater environments in continental Southeast Asia (*Amyda ornata*, *Cuora couro*, *Cyclemys oldhamii*, *Heosemys annandalii*, *Heosemys grandis*, and *Siebenrockiella crassicollis*) in Cambodia (Platt et al., 2008), Vietnam (Le, 2007), and Laos (Brakels, 2018).

### Biodiversity Changes Information Provided by the Studied Archaeological Samples

The taxa identified at the investigated sites provide information about the local changes of turtle fauna across the Holocene. These can be categorized into species that were: I) not or weakly impacted, II) species whose distribution shrank, and III) species that went extinct.

#### I) Weakly Impacted Species (N=7)

*Manouria impressa* is still present today in the two areas where it has been identified (northern Thailand and southwest Cambodia) (Rhodin et al., 2021) (Figure 11-A). This turtle is a terrestrial species living in mountainous forests, mostly active during the hot and rainy season, and primarily consumes mushrooms (Wanchai et al., 2013; Cota et al., 2021). Regarding its occurrence in Laang Spean, it is interesting to note that the site is not in an elevated area, which could indicate a past occurrence of *M. impressa* in low-elevation forests that no longer exist nowadays, but the data are currently too scarce to investigate this question. The same remarks could be made about *Manouria emys* (Claude et al., 2019). The species of *Heosemys* identified in the samples are all present in the vicinity of the sites today (Figure 11-B). *Heosemys grandis* and *H. annandalii* are freshwater species commonly found in swamps, while *H. spinosa* is a lowland forest species (Sharma & Tisen, 2000; Das, 2010). Some other taxa might have been impacted but only in part of their range. For instance, *Cuora couro* (Figure 11-C) and *Malayemys* sp. (Figure 11-D) are both still present today in the areas of the sites where they have been identified but not in Laang Spean. However, both of these taxa are still present a few dozen kilometers away from this last site, which indicates that if their distribution area has effectively shrunk, this might be a very recent phenomenon. *Cuora couro* is a semi-aquatic species inhabiting many types of lowland freshwater habitats. It is mostly active at night and can feed both in water and on land (Schoppe & Das, 2015). *Malayemys* is a genus of aquatic species that estivate on land in forests and occupy a variety of freshwater habitats (Dawson et al., 2018, 2020). Regarding Trionychidae, we identified *Dogania* inside its modern range in Southern Thailand (Fig. 12-B).

**Figure. 11.**
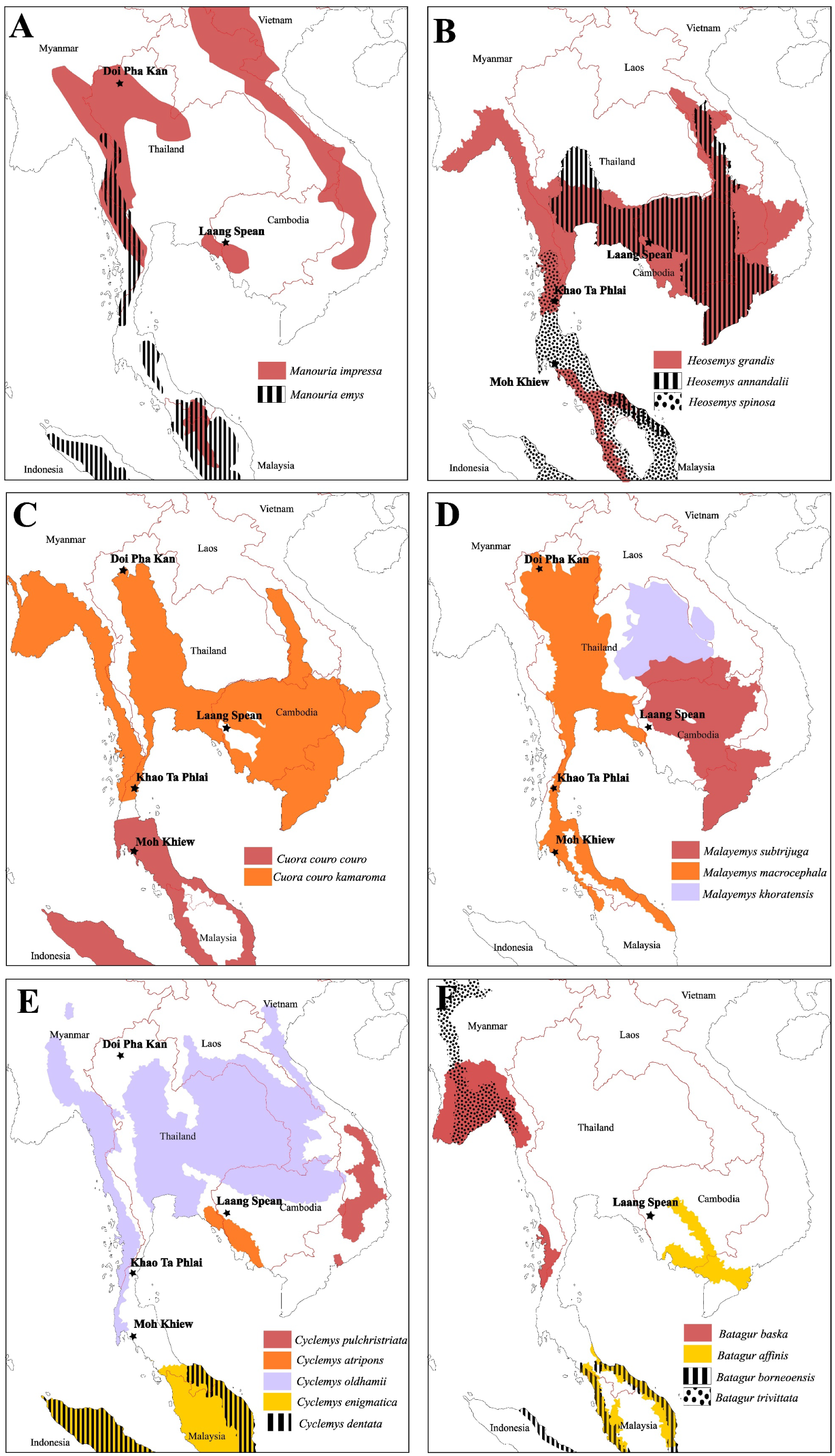
Modern distribution maps of the Testudinidae and Geoemydidae species identified in the four archaeological assemblages, along with the locations of the archaeological sites that provided remains of the concerned taxa. The distribution data are from Rhodin et al. (2021) and Chaianunporn et al., (2025) for *Malayemys*.

**Figure. 12.**
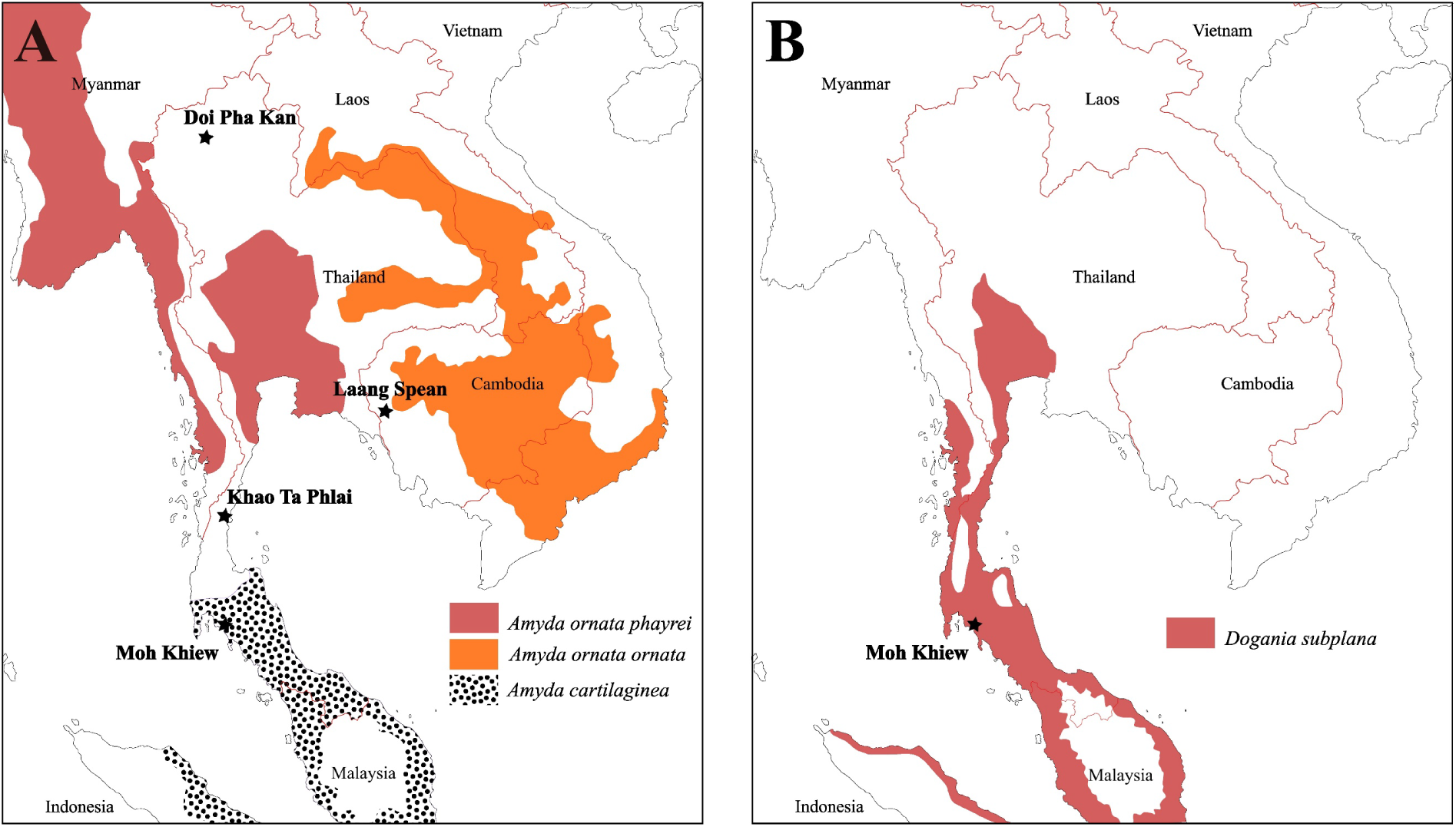
Modern distribution maps of the Trionychidae species identified in the four archaeological assemblages, along with the locations of the archaeological sites that provided remains of the concerned taxa. The distribution data are from Rhodin et al. (2021).

#### II) Species Whose Distribution Area Shrunk (N=4)

The distribution of *Cyclemys* cf. *oldhamii* may have undergone significant changes, as it is now absent in the vicinity of all the sites, although it has been recorded a few dozen kilometers away from Laang Spean, Khao Ta Phlai, and Moh Khiew (Figure 11-E). This could reflect either shrink of distribution, lack of survey of modern species, or transportation by prehistoric populations. However, in Doi Pha Kan, the distance to the modern distribution area is around 100 km, making the hypothesis of a reduction in the distribution area of this taxon more likely although it might still be related to a lack of biodiversity survey in that area. A similar observation can be made for our finding of *Batagur* in Laang Spean (Figure 11-F). *Batagur affinis* (Cantor 1847) is the only species of the *Batagur* genus currently present in Cambodia. It inhabits the floodplain of the Tonle Sap, which is only 50 km away from Laang Spean (Moll et al., 2015). It is an aquatic species living in estuaries and tidal regions of large rivers (Moll et al., 2015; Thomson, 2021). The distribution of these two species has undoubtedly shrunk since the early/middle Holocene. *Indotestudo elongata*, previously mentioned in Laang Spean (Bochaton et al., 2023) but not described in this paper, should also be added to this list, as its closest modern occurrence is around 50 km away from the site. Regarding the softshell turtle *Amyda* (Fig. 12-A), we identified its remains in Doi Pha Kan in Northern Thailand, over 150 km from its closest modern range near the Myanmar border (Rhodin et al., 2021) but, although this genus might be rare and absent in several areas of northern Thailand, its complete absence could reflect a lack of documentation in several areas (J. C. pers. obs). *Amyda* was also identified in Khao Ta Phlai between the modern distributions of *A. ornata* and *A. cartilaginea* (Fig. 12) in an area where the genus is absent nowadays. Bones of this genus have also been found at the Laang Spean site, although *Amyda* is still present just a few kilometers from this site today. Based on its modern range, it is likely that *Amyda* has experienced a significant reduction in its distribution area in Northern Thailand.

#### III) Species That Went Extinct (N=1 or 2)

The only clear extinction, or at least a massive reduction in distribution, is associated with the one or two species of Testudinidae (*Geochelone* sp. and Testudinidae indet.) identified in the archaeological samples. Whether the material is attributed to *Geochelone* or another Testudinidae taxon, it indicates that at least one species of this family was present in Thailand and Cambodia in the past and that it completely disappeared over the course of the Holocene. Such event is unlikely to reflect an overexploitation by human groups as the remains of these turtles are very scarce in the archaeological assemblages. There is currently not a single extinction of turtle species documented in Southeast Asia during the Late Pleistocene and the Holocene (Rhodin et al., 2015).

### Has Southeast Asian Turtle Biodiversity Changed in the Last 10,000 Years?

The scarcity of available data regarding past turtle biodiversity in Southeast Asia currently precludes detailed analyses, but some interesting observations can be made. Our results clearly indicate that the different investigated areas had varying trajectories concerning the evolution of their turtle biodiversity, as the impacted or non-impacted species were not randomly distributed across the different sites. The two sites in southern Thailand show a turtle fauna that was very close to the modern one, with only one species that may have suffered a limited reduction in its distribution range (*Cyclemys* sp.). The situation is similar in northern Thailand (Doi Pha Kan), even though Trionychidae turtles seem to have disappeared from that area, and an unknown extinct testudinid species is present in this deposit. It is currently impossible to link this extinction or change of distribution to a specific phenomenon, as it is presently impossible to estimate when this species disappeared. Aside from this observation, the turtle fauna of Thailand appears to exhibit remarkable stability, which stands in stark contrast to the observations made in the central plain of Thailand (Claude et al., 2019). In this latter region, at least two forest taxa and one large freshwater turtle may have been extirpated following environmental changes and deforestation in the second half of the Holocene. This suggests that not all areas of Thailand were impacted in the same way. Regarding the sites we studied, the only locality that truly stands out in our sample is Laang Spean, where seven of the eleven species we identified (Table. 01 – *Batagur* sp., *Cuora couro*, *Cyclemys* sp., *Geochelone* sp., *Indotestudo elongata*, and *Amyda* sp.) are absent from the surrounding area (Figs. 11 and 12).

We believe that the biodiversity modifications observed at the sites reflect recent human impact rather than deep historical alterations of the environments, and this is for two reasons: 1) the distribution changes observed are mostly minor, typically only a few dozen kilometers in most cases, and 2) the fact that the Cambodian site shows the most significant changes in its turtle fauna compared to the sites in Thailand can be easily explained by the magnitude of the very recent environmental modifications these two countries have experienced. Indeed, compared to Thailand, the impact of deforestation has been much more severe in Cambodia. Thailand’s forest cover was estimated to be about 70% of the country’s surface in 1930 (35.7 million ha), shrinking to around 29.6 million ha in 1952 and about 17.8 million ha in 1978 (Ramitanondh, 1989). This corresponds to an annual deforestation rate of r=1.4%. However, the deforestation rate has significantly decreased since then, being reduced by more than tenfold between 1990 and 2016, and maintaining an annual rate of r=0.028% until at least 2020 (Puyravaud, 2003; FAO, 2020a; b). In Cambodia, the situation is very different. Forest cover represented 73.3% of the country’s area in the 1960s (13.3 million ha) but decreased to 10.937 million ha by 1990 (Tsujino et al., 2019; FAO, 2020c), corresponding to an annual deforestation rate of r=0.6%. The subsequently documented annual deforestation rate was r=0.23% between 1990 and 2010, but this rate dramatically increased to r=4.5% between 2010 and 2015, and r=2.1% between 2015 and 2020 (Puyravaud, 2003; FAO, 2020b; c). This situation is already well known, and the future of Cambodia’s biodiversity is now the most concerning among continental Southeast Asian countries (Botterill-James et al., 2024). Laang Spean, being located outside of a protected area, has suffered massive and very fast environmental changes in recent decades. This observation, if not related to an overall paucity of biodiversity inventory in Cambodia, could be a striking demonstration of the quick effect of massive environmental alterations that could wipe out taxa proven to be strongly resilient even in disturbed areas of Thailand.

Overall, our data point to the resilience of turtle assemblages outside the areas that have been heavily anthropized, such as the central plain of Thailand and Cambodia. In these two regions, the extent of environmental modifications has exceeded the adaptation capacity of the taxa, leading to a significant reduction in local biodiversity. That said, other factors might also be at play, such as climatic events and human-induced changes in the distant past. These phenomena are characterized by significant reductions in geographic distribution and even the extinction of some rare taxa, but these events cannot be accurately dated and thus characterized at this time. However, these impacts appear to be limited, as most local species assemblages have persisted since the Late Pleistocene/Early Holocene without major modifications. This could also be the case in other Southeast Asian area. For instance, in the Late Pleistocene archaeological layers of Niah Cave in Sarawak, the three species identified in the second phase and the three species identified in the first phase are still extant in the vicinity of the site today (Pritchard et al., 2009). Only the study of additional bone assemblages and well-dated occurrences of critically important taxa will enhance our understanding of the causes behind these modifications.

## Conclusion

Our observations indicate that the turtle assemblages in northern and southern Thailand, and southwest Cambodia, have experienced only limited changes across the Holocene, with the exception of Laang Spean in Cambodia. We also raise questions about the existence of one or several potentially extinct species of Testudinidae in continental Southeast Asia at an unknown time during the Holocene. Additionally, we observed significant reductions in geographic distribution affecting some species, such as *Batagur*, *Cyclemys*, and *Amyda*. Unfortunately, it is currently difficult to determine the timing and causes of the biodiversity changes we observed, and we have had to rely on probabilistic hypotheses. To address these uncertainties, it is now necessary to improve the quality of the available paleontological and zooarchaeological records, particularly for more recent periods (historical and modern). Radiometric dating of the remains of critically important taxa will be essential to clarify their periods of occurrence in different areas. This should be done in parallel with the enhancement of modern reference collections and possibly molecular approaches to identify past taxonomic diversity in Southeast Asia as reliably as possible. In conclusion, although the results described here show that it is not too late to preserve the native biodiversity of several areas in continental Southeast Asia, our observations serve as a warning of the potential fate of turtle biodiversity in this region if further loss of natural environments, such as what is currently happening in Cambodia, is not prevented. We also emphasize the importance of biodiversity surveys to accurately track the evolutionary dynamics of species facing environmental disturbances.

## Funding

This work has been done thanks to the support of several funding agencies: the FYSSEN foundation, the DIM-MAP funded by the Region île-de-France, the French Ministry of Europe and Foreign Affairs, and the IRN PalBioDivASE 0846 funded by the CNRS. It was also supported by the Mission Préhistorique FrancoCambodgienne, the Mission Préhistorique Franco-Thaïe, and the Mission Paléolithique Franco-Thaïe of the French Ministry of Europe and Foreign Affairs (MEAE, Paris).

## Conflict of interest disclosure

The authors declare that they comply with the PCI rule of having no financial conflicts of interest in relation to the content of the article.

## Author contributions

**Conceptualization**: Corentin Bochaton; **Data curation**: Corentin Bochaton; **Formal analysis**: Corentin Bochaton, Julien Claude; **Funding acquisition** : Corentin Bochaton, Valéry Zeitoun, Hubert Forestier; **Investigation** : Corentin Bochaton, Wilailuck Naksri, Sirikanya Chantasri, Melada Maneechote, Supalak Mithong, Julien Claude; **Methodology** : Corentin Bochaton; **Resources** : Sophady Heng, Hubert Forestier, Prasit Auetrakulvit, Valéry Zeitoun, Jutinach Bowonsachoti; **Supervision** : Corentin Bochaton; **Visualization**: Corentin Bochaton; **Writing – original draft**: Corentin Bochaton, Julien Claude; **Writing – review & editing**: Corentin Bochaton, Julien Claude.

## Appendix 1

List of the specimens used in the study. <u>Abbreviations</u>: **CUMZ-R**: Chulalongkorn University Museum of Zoology (Bangkok, Thailand) reptile collection; **MNHN-RA**: Muséum national d’Histoire naturelle (Paris, France) reptiles and amphibians collection; **MNHN-RA**: Muséum national d’Histoire naturelle (Paris, France) comparative anatomy collection; **MRHNB**: Musée Royal d’Histoire naturelle de Belgique; **MVZ**: Museum of Vertebrate Zoology (Berkeley, USA); **UF**: University of Florida Museum of Natural History (Gainesville, USA).

**Geoemydidae:** *Batagur baska* (MNHN-RA-0.9094; CUMZ-R-TG1105; CUMZ-R-TG1108; CUMZ-R-TG151697; CUMZ-R-TG155191); *Cuora couro* (MNHN-RA-1889-182; CUMZ-R-TG364; CUMZ-R-TG342; UF 134824; UF 134833); *Cuora galbinifrons* (UF 81775; UF 154221); *Cuora mouhotii* (UF 53668; UF 61986; UF 81980); *Cuora trifasciata* (UF 103395); *Cyclemys oldhamii* (CUMZ-R-TG969); *Cyclemys atripons* (UF 105993; UF 119584, UF 116449); *Cyclemys dentata* (CUMZ-R-TG954; UF 135096; UF 139828; UF 154136; UF 154138; UF 155192); *Cyclemys* sp. (CUMZ-R-TG959; CUMZ-R-TG960; CUMZ-R-TG1432; CUMZ-R-TG1053); *Heosemys annandalii* (CUMZ-R-TG606; CUMZ-R-TG608; CUMZ-R-TG609; CUMZ-R-TG613; CUMZ-R-TG614; CUMZ- R-TG615; CUMZ-R-TG618; CUMZ-R-TG619; CUMZ-R-TG622; CUMZ-R-TG624; CUMZ-R- TG633; CUMZ-R-TG647; CUMZ-R-TG656; CUMZ-R-TG762; UF 154245; UF 155196); *Heosemys grandis* (CUMZ-R-TG863; UF 154202; UF 154244); *Heosemys spinosa* (CUMZ-R-TG1005; UF 32853; UF 154230; UF 154227); *Leucocephalon yuwonoi* (UF 97335; UF 109835); *Malayemys subtrijuga* (CUMZ-R-TG42; CUMZ-R-TG51; CUMZ-R-TG58; CUMZ-R-TG147; MNHN-RA- 1874-78; UF 62085; UF 67527; UF 67638; UF 69380; UF 150281); *Mauremys annamensis* (UF 88811; UF 151054; UF 154218); *Mauremys mutica* (UF 107174; UF 88806); *Melanochelys trijuga* (CUMZ-R-TG1151; MNHN-RA-1900-184; UF 61987); *Morenia ocellata* (MNHN-RA-0.9167); *Notochelys platynota* (UF 123862; UF 151154; UF 154260); *Orlitia borneensis* (MRHNB 1941.97; UF 152476; UF 154020; UF 154234); *Siebenrockiella crassicollis* (CUMZ-R-TG551; MNHN-ZM-AC 1875-88). **Testudinidae**: *Geochelone elegans* (MVZ-Herp-23519); *Geochelone platynota* (AMNH-R-57451); *Manouria emys* (CUMZ-R-TT11; CUMZ-R-TT12; CUMZ-R-TT15; UF 52644; UF 27470; UF 67618; UF 134830; UF 155203). **Platysternidae**: *Platysternon megacephalum* (UF 56669; UF 56950; UF 57111; UF 57608). **Trionychidae**: *Amyda cartilaginea* (UF 57728); *Dogania subplana* (UF 61810; UF 69259); *Lissemys punctata* (UF 56017); *Pelodiscus sinensis* (UF 175673; UF 175674; UF 175673; UF 175674).

## Appendix 2

Supplementary figures: Drawings of the carapace and plastron of the modern Southeast Asian turtles (Testudinidae and Geoemydidae). The species are ordered alphabetically and *Indotestudo elongata* is not included as already figured in Bochaton et al. (2023).

---

**Fig. S1.**
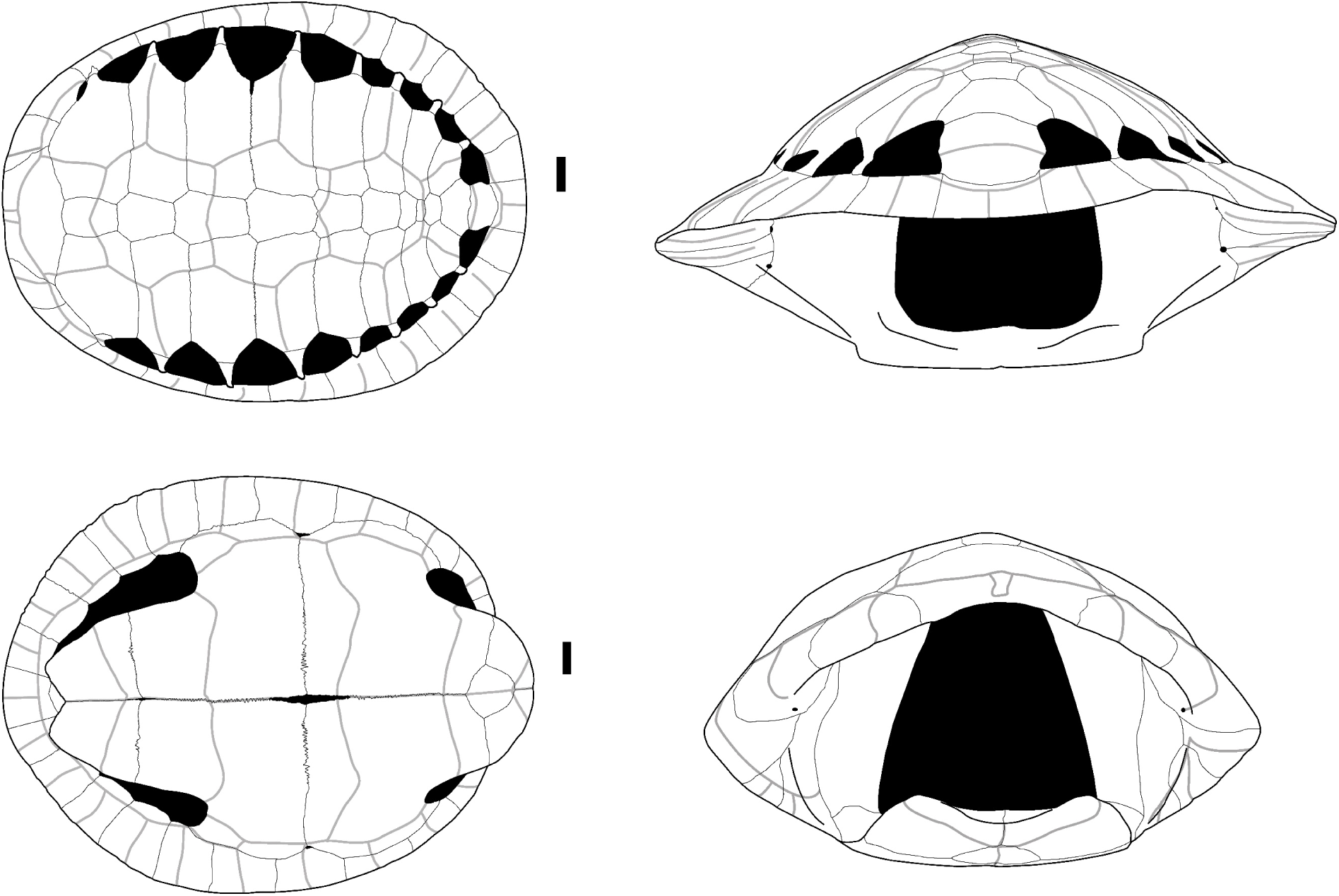
Carapace and plastron of a subadult specimen of *Batagur borneoensis* (Schlegel & Müller, 1845) (CUMZ-R-TG1108) in dorsal view (upper left), ventral view (lower left), posterior view (upper right), and anterior view (lower left). Scalebars=2cm.

**Fig. S2.**
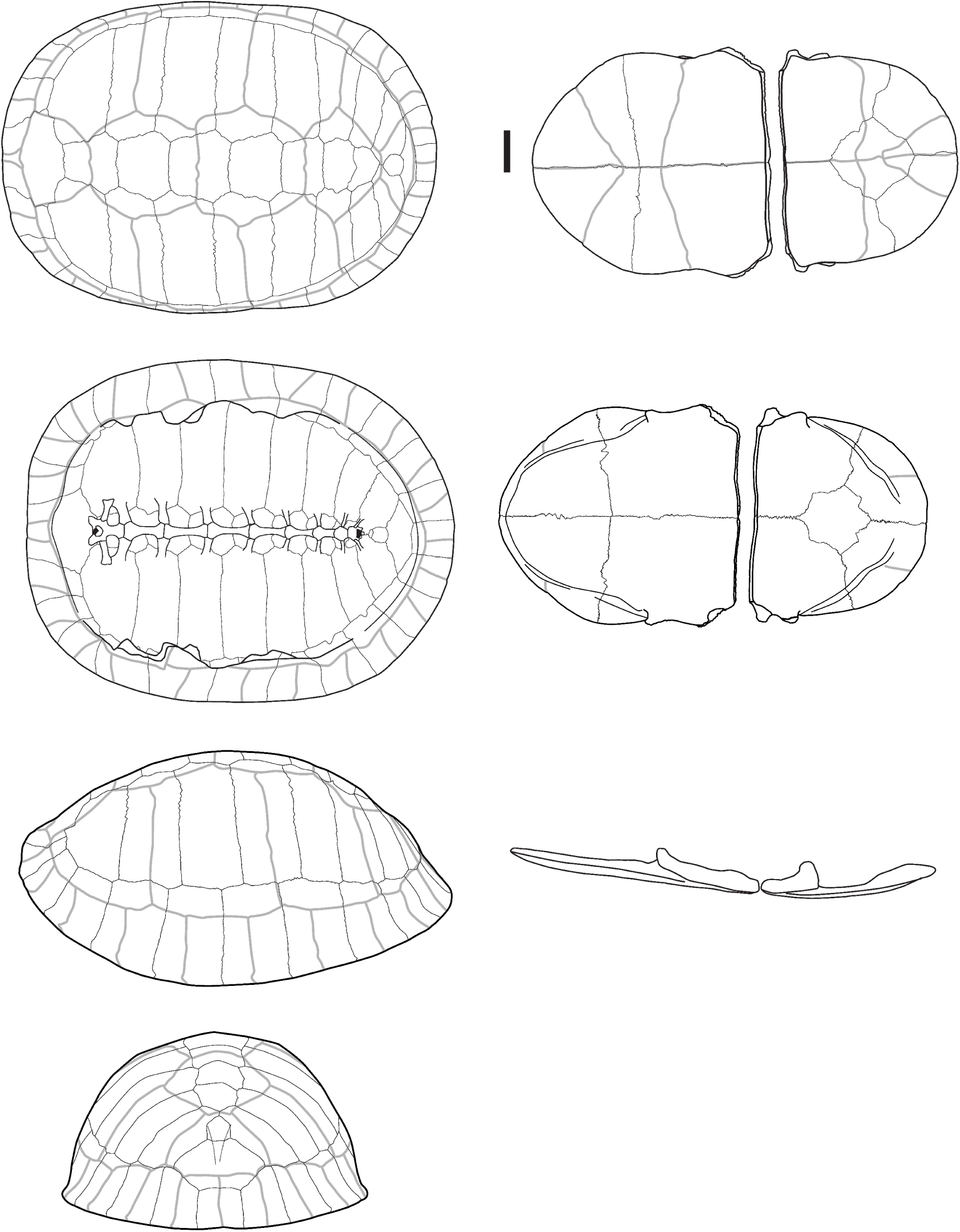
Carapace (left) and plastron (right) of *Cuora couro* (Leschenault de la Tour in Schweigger, 1812) (CUMZ-R-TG342). From top to bottom for carapace: dorsal, ventral, lateral, and posterior views, and for plastron: ventral, dorsal, and lateral views. Scalebar=2cm.

**Fig. S3.**
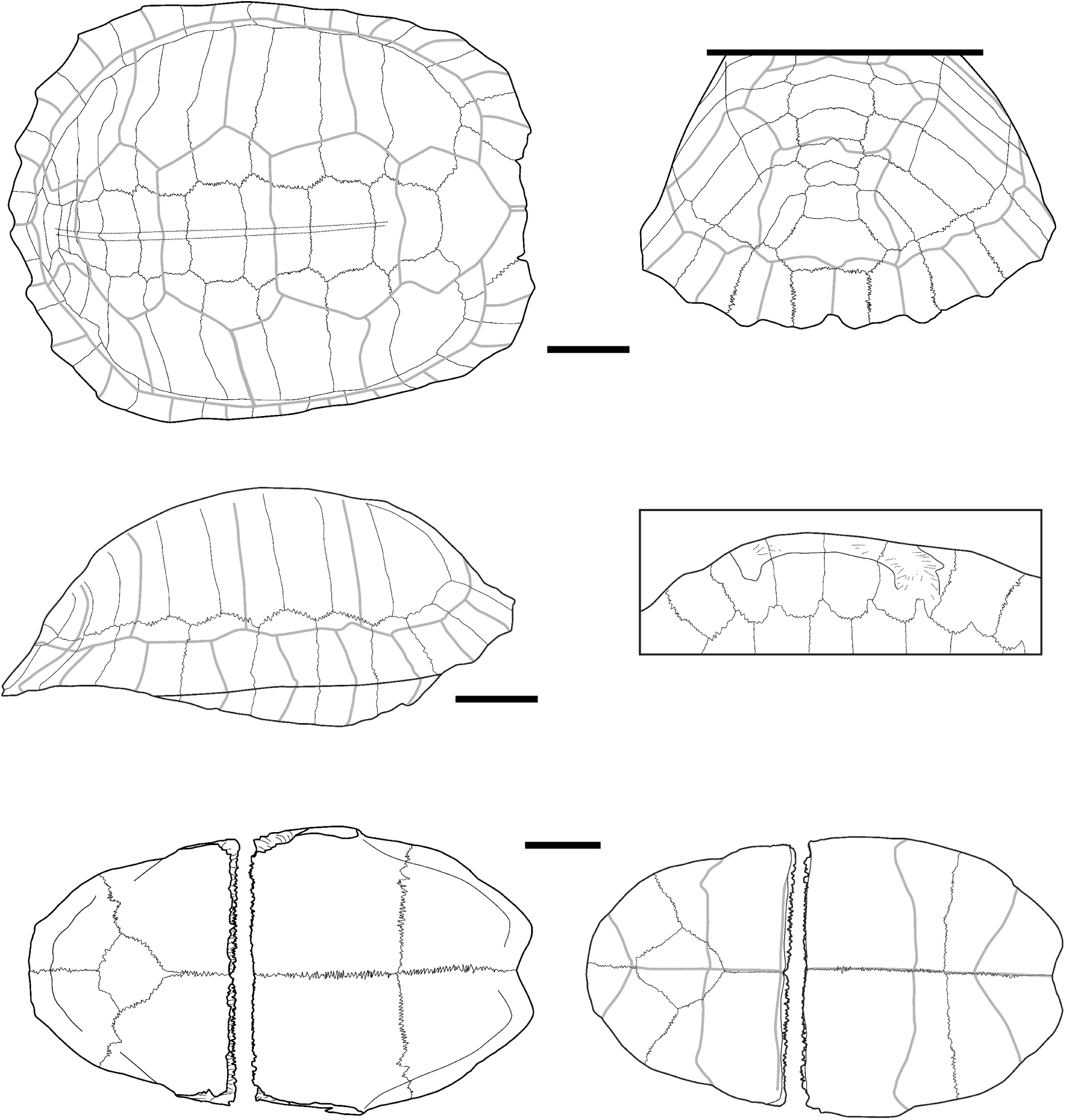
Carapace and plastron of *Cuora mouhotii* (Gray, 1862) (UF 81980). Carapace in dorsal (upper left), posterior (upper right), and lateral (medium left) views. Plastron in dorsal (bottom left) and ventral (bottom right) views. A detailed view of the attachment surface of the carapace with the plastron is also figured (medium right). Scalebars=2cm.

**Fig. S4.**
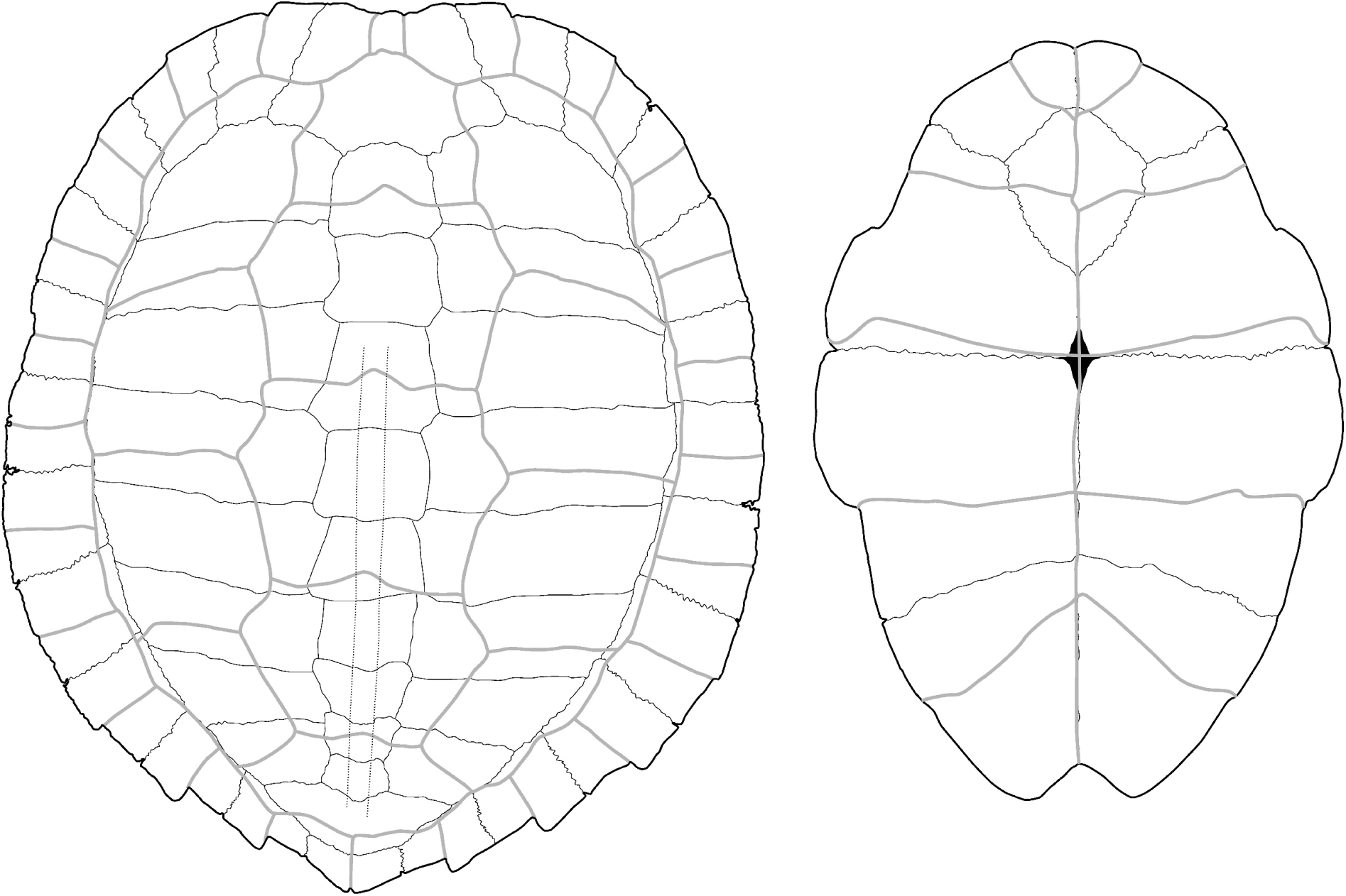
Carapace (left) and plastron (right) of *Cyclemys atripons* Iverson & McCord, 1997 (UF 116449). No scalebar.

**Fig. S5.**
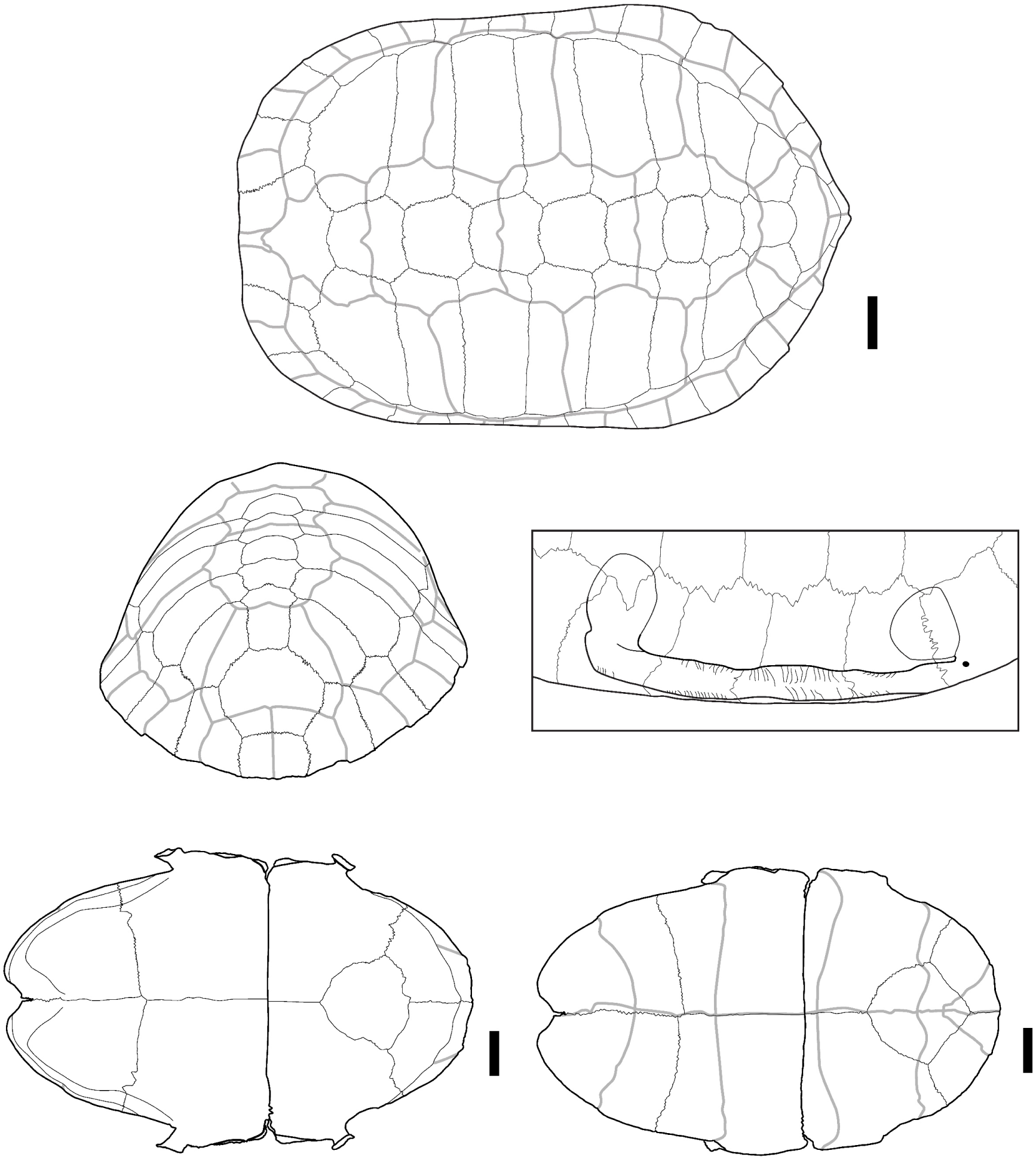
Carapace and plastron of *Cyclemys oldhamii* Gray, 1863 (CUMZ-R-TG969). Carapace in dorsal (upper), and posterior (medium left) views. Plastron in dorsal (bottom left) and ventral (bottom right) views. A detailed view of the attachment surface of the carapace with the plastron is also figured (medium right). Scalebars=2cm.

**Fig. S6.**
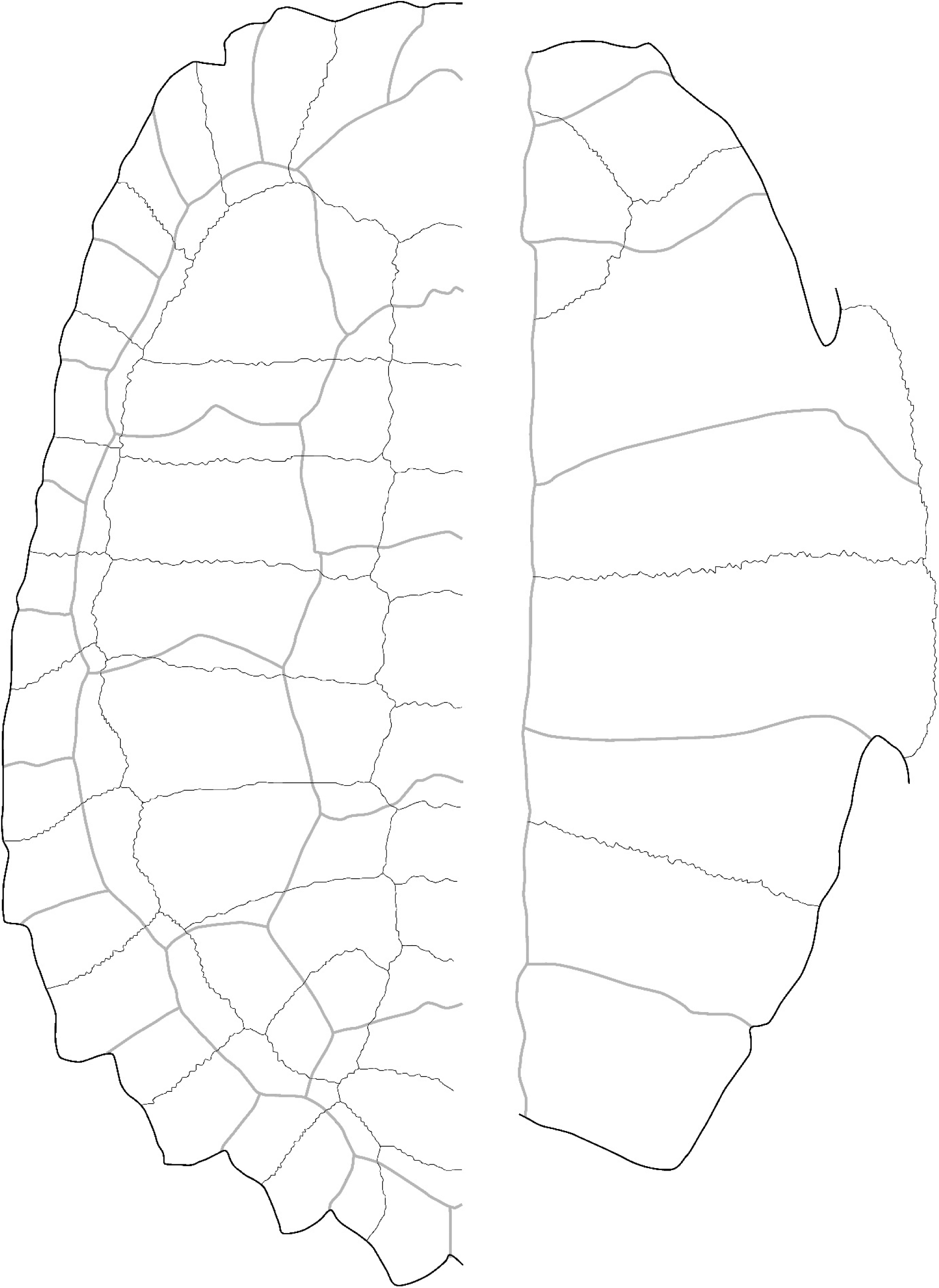
Carapace (left, dorsal view) and plastron (right, ventral) of *Leucocephalon spengleri* (Gmelin, 1789) (PCHP 12207). Modified from Ascarrunz, Claude, and Joyce (2021). No scalebar.

**Fig. S7.**
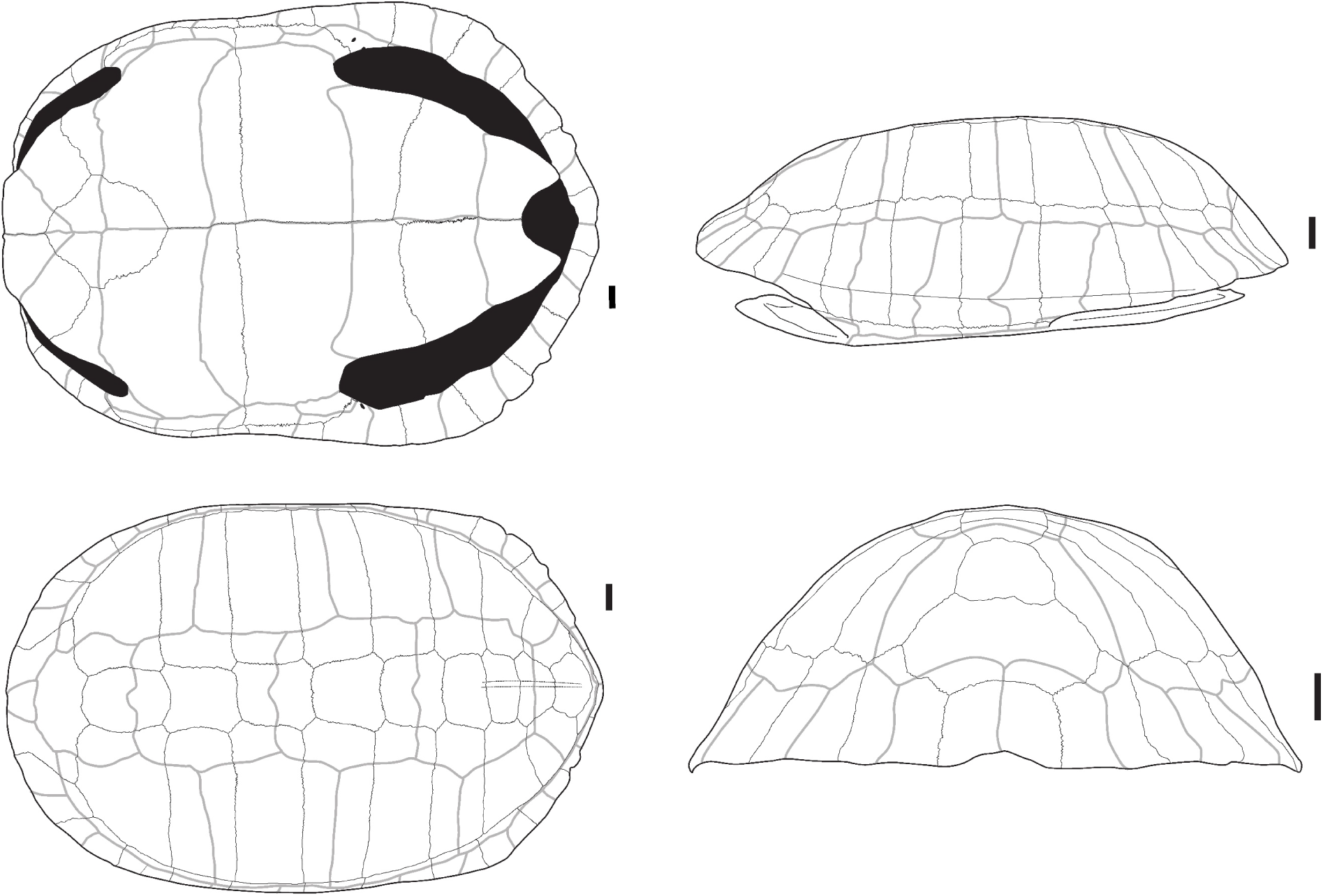
Carapace of *Heosemys annandalii* (Boulenger, 1903) (CUMZ-R-TG647) in ventral (upper left), dorsal (bottom left), lateral (upper right), and posterior (bottom right) views. Scalebars=2cm.

**Fig. S8.**
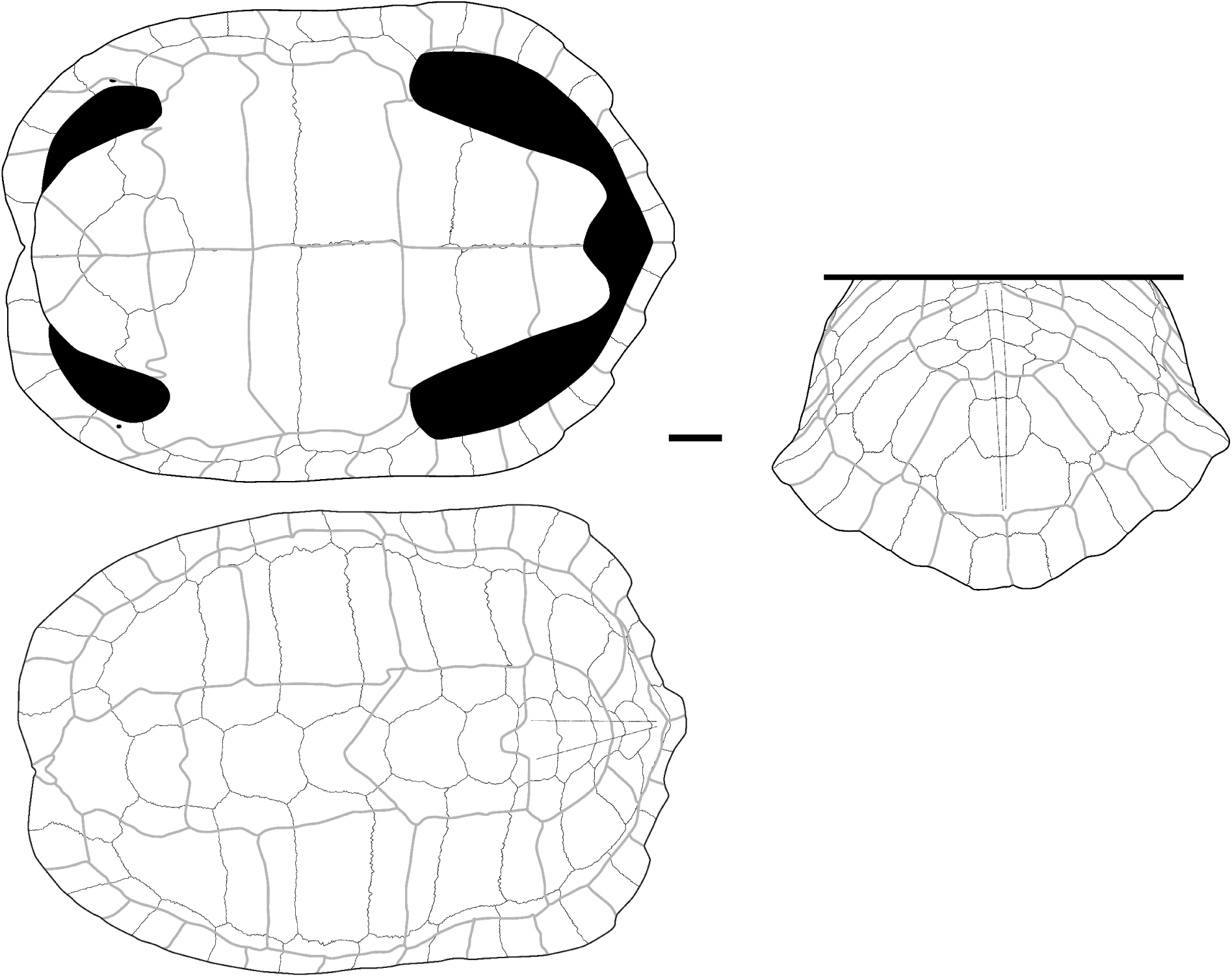
Carapace of *Heosemys grandis* (Gray, 1860) (Chulalongkorn Museum – no number) in ventral (upper left), dorsal (bottom left), and posterior (right) views. Scalebar=4cm.

**Fig. S9.**
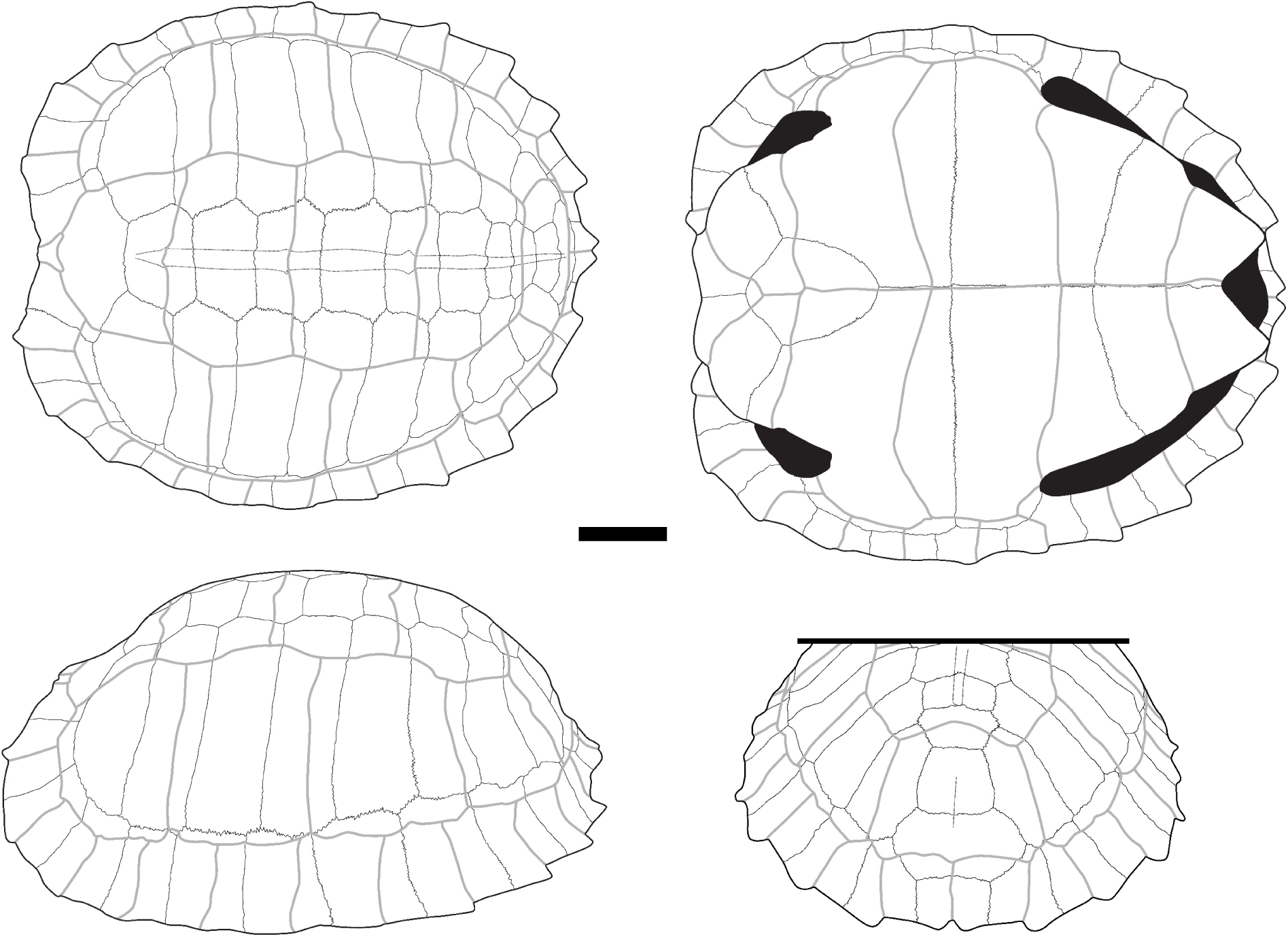
Carapace of *Heosemys spinosa* (Gray, 1830) (UF 32853) in dorsal (upper left), ventral (upper right), lateral (bottom left), and posterior (bottom right) views. Scalebar=3cm.

**Fig. S10.**
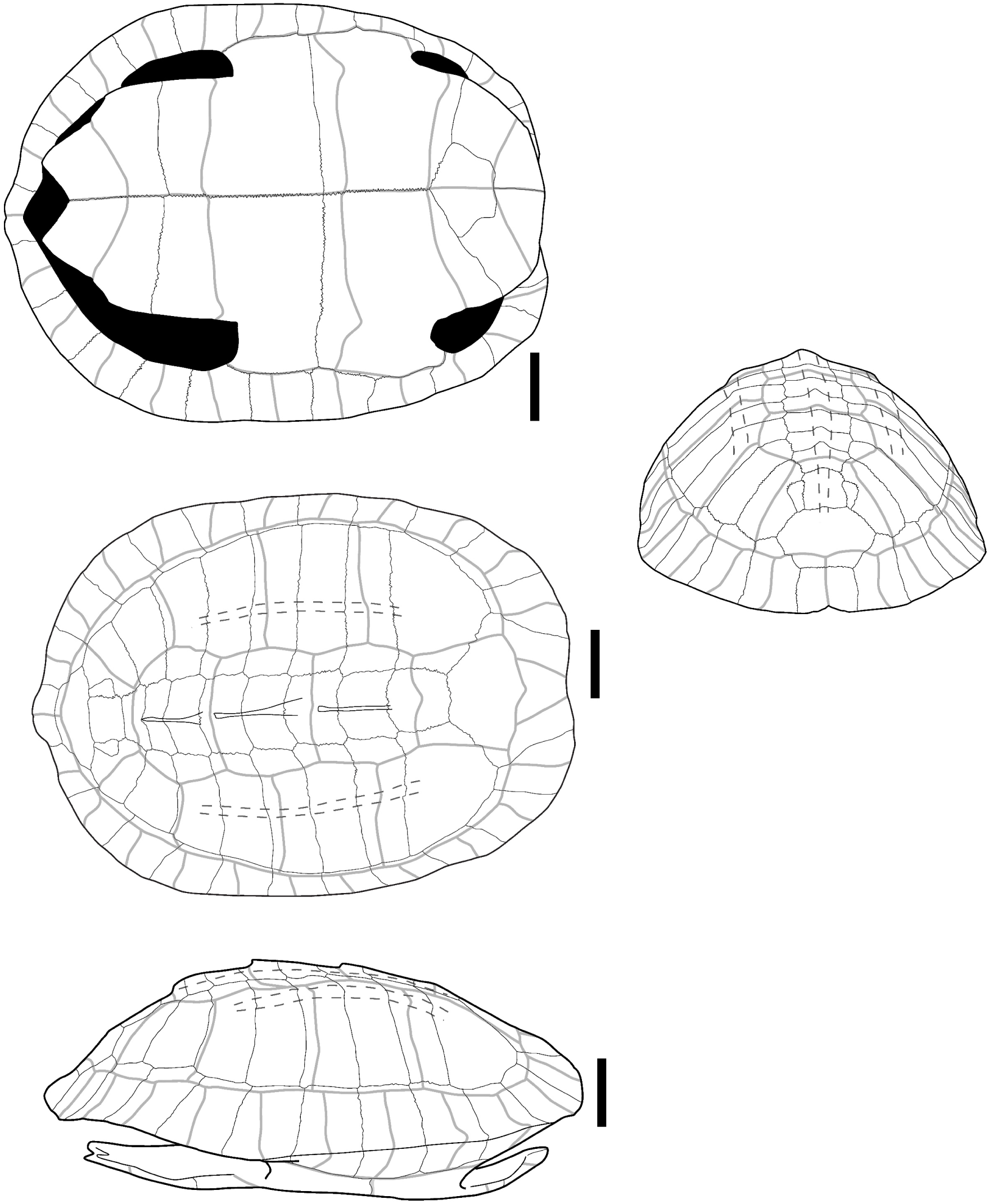
Carapace of *Malayemys macrocephala* (Gray, 1859) (CUMZ-R-TG42) in ventral (upper left), dorsal (medium left), lateral (bottom left), and posterior (right) views. Scalebar=2cm.

**Fig. S11.**
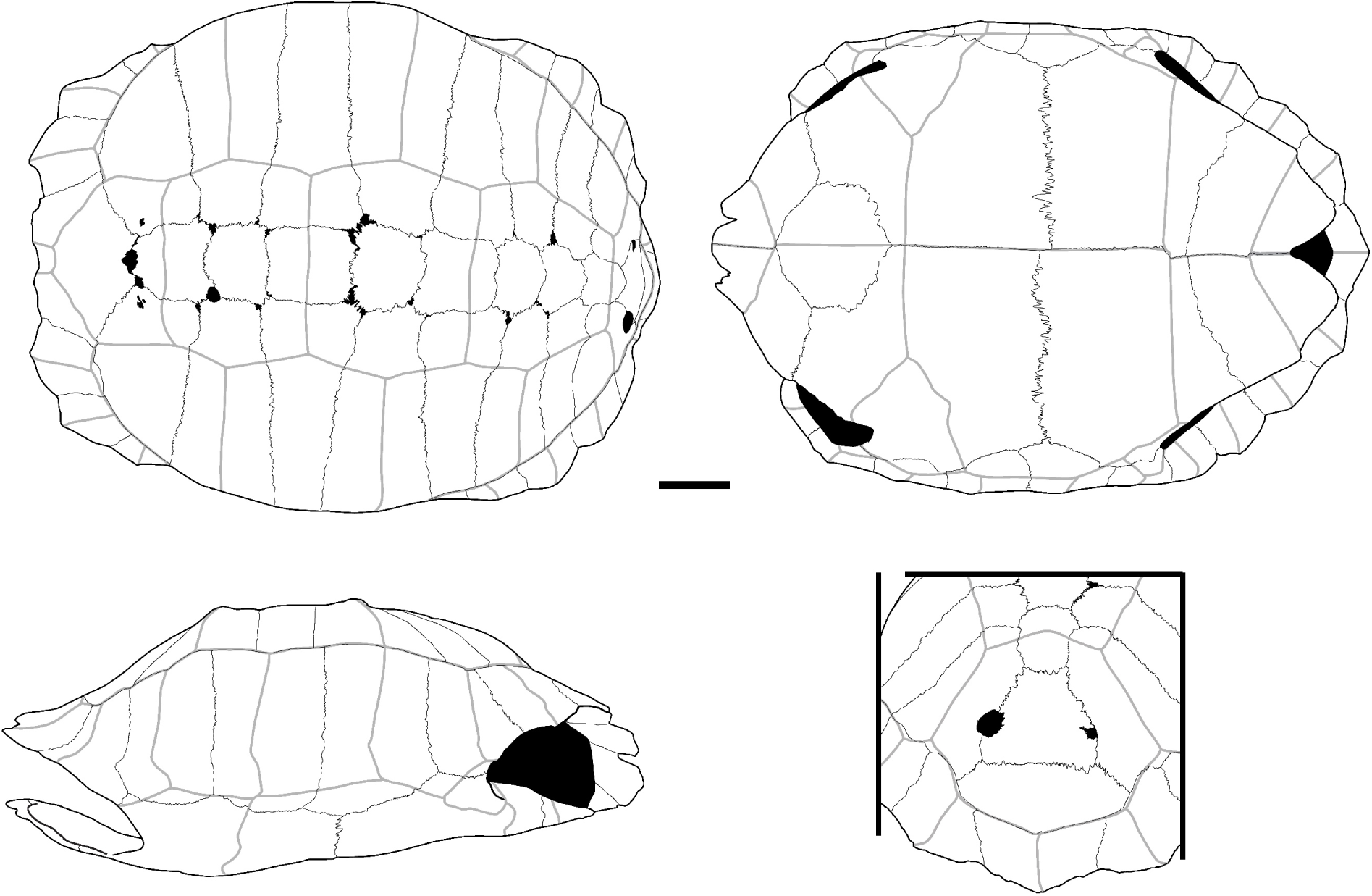
Carapace of *Manouria emys* (Schlegel & Müller, 1840) (UF 134830) in dorsal (upper left), ventral (upper right), lateral (bottom left), and posterior (bottom right) views. Scalebar=5cm.

**Fig. S12.**
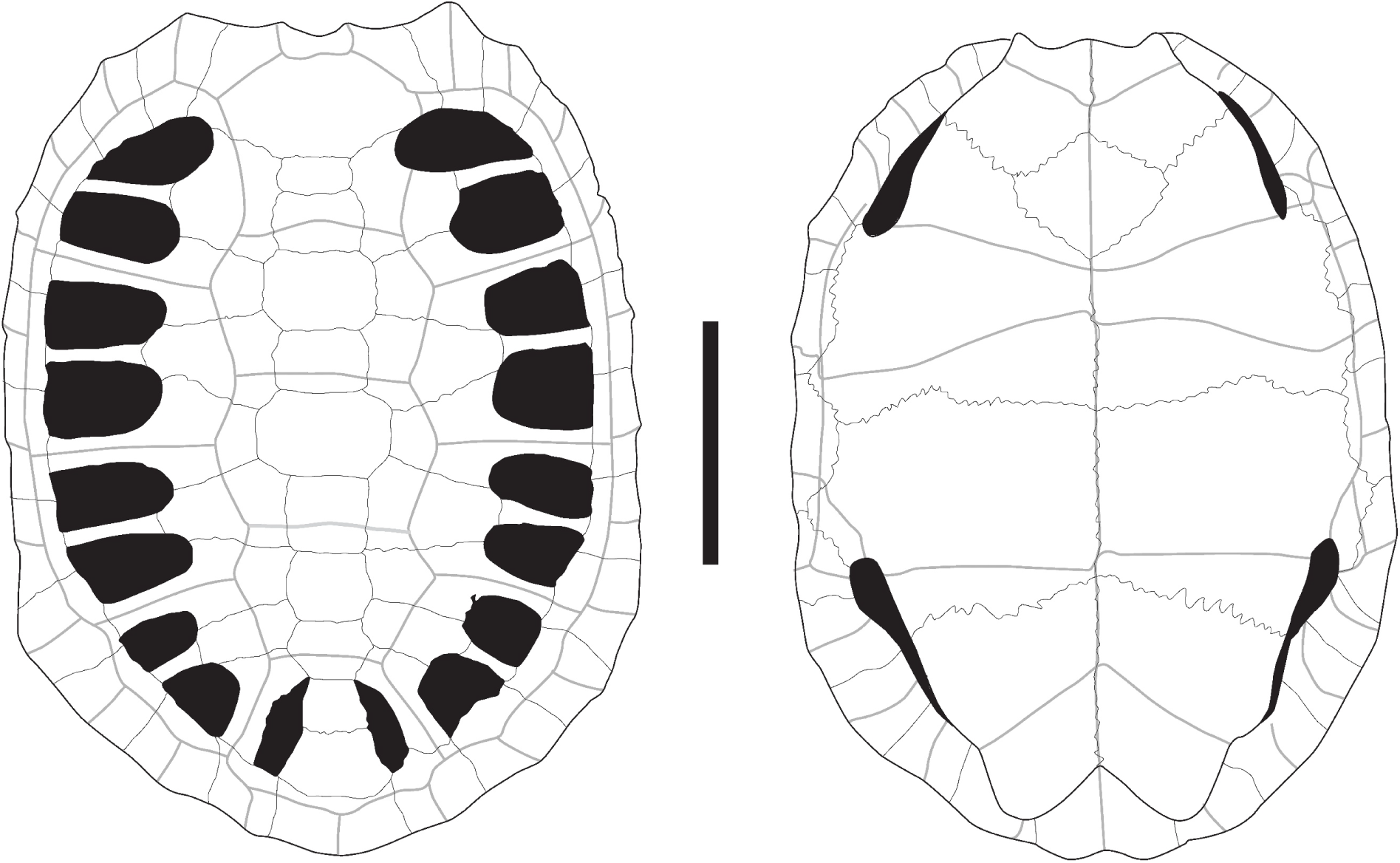
Carapace of *Manouria impressa* (Günther, 1882) modified from Bourret (1941) in dorsal (left) and ventral (right) views. Scalebar=5cm.

**Fig. S13.**
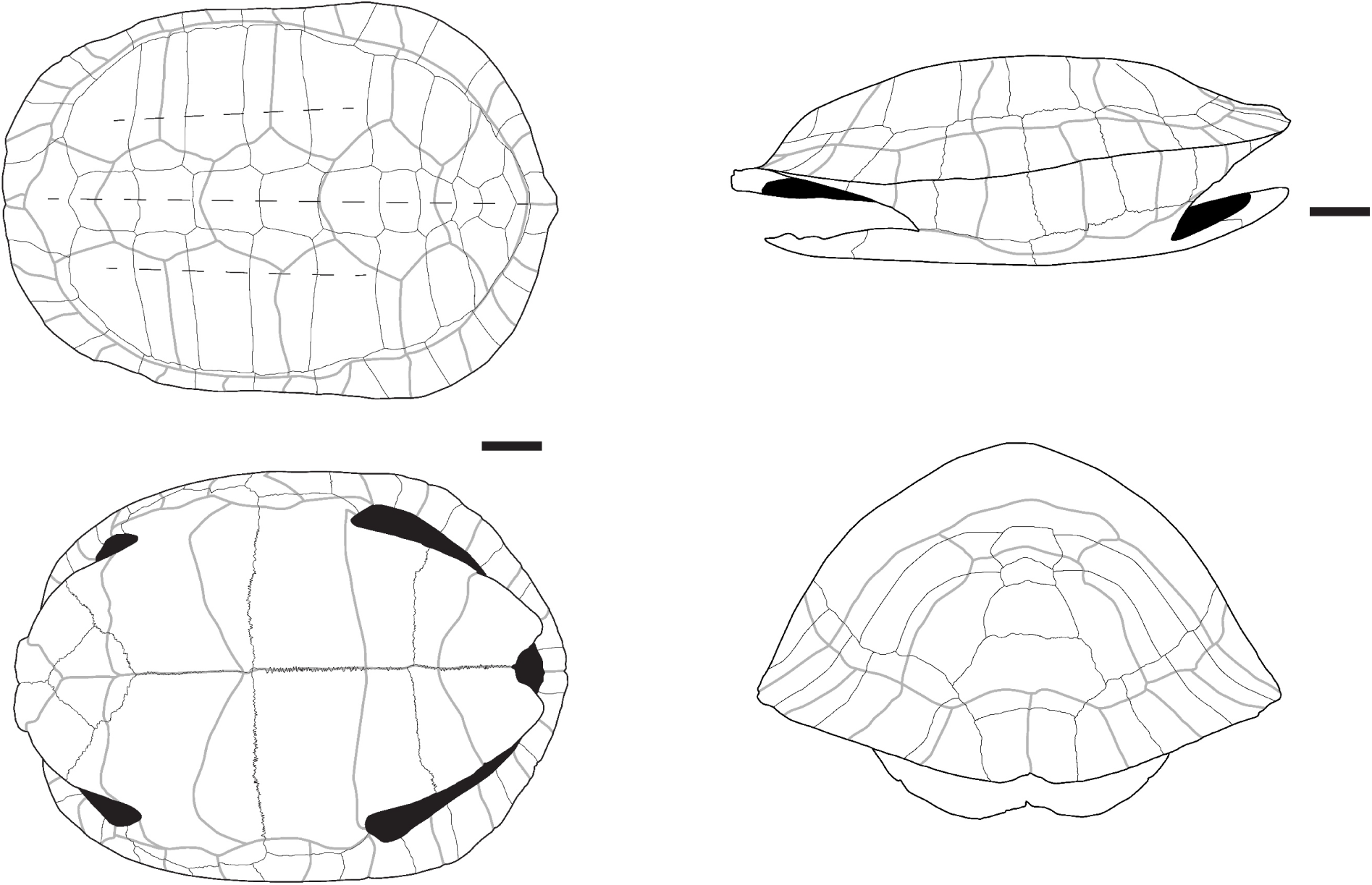
Carapace of *Melanochelys trijuga* (Schweigger, 1812) (CUMZ-R-TG51) in dorsal (upper left), ventral (lower left), lateral (upper right), and posterior (bottom right) views. Scalebars=3cm.

**Fig. S14.**
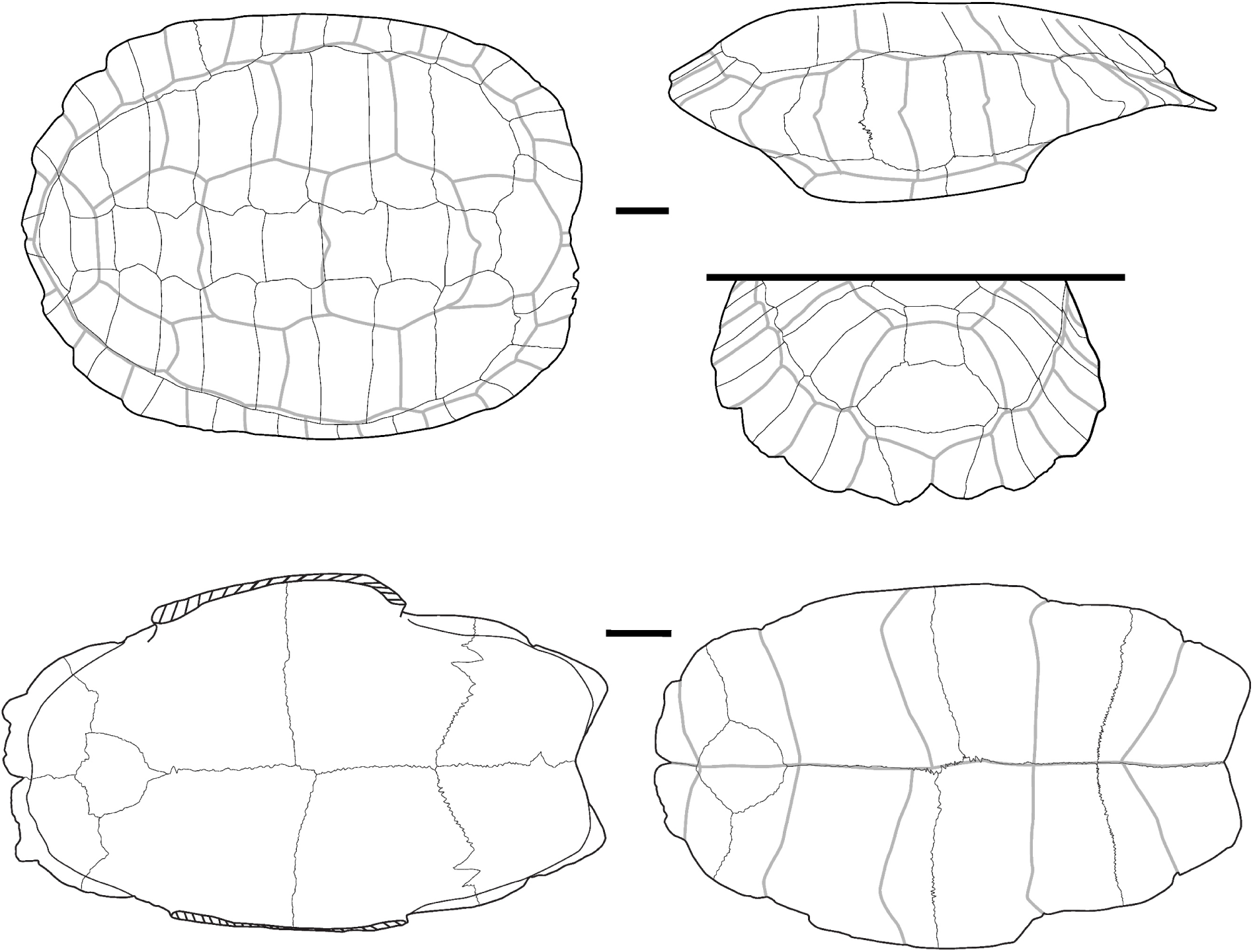
Carapace and plastron of *Notochelys platynota* (Gray, 1834) (UF 123862). Carapace in dorsal (upper left), lateral (upper right), and posterior (medium right) views. Plastron in dorsal (bottom left) and ventral (bottom right) view. Scalebars=2cm.

**Fig. S15.**
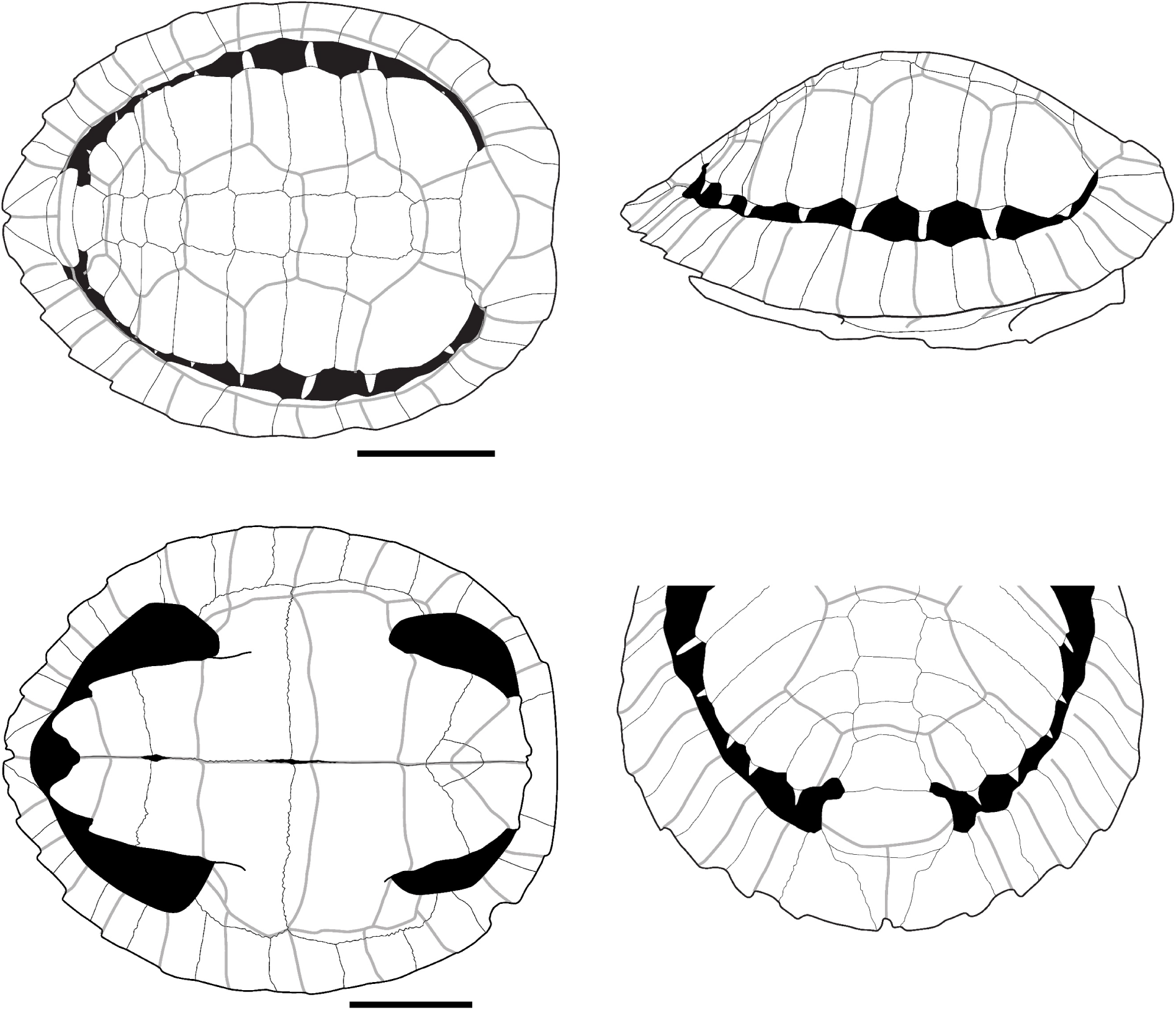
Carapace of a juvenile specimen of *Orlitia borneensis* Gray, 1873 (MRHNB 1941.97) in dorsal (upper left), ventral (bottom left), lateral (upper right), and posterior (bottom right) views. Scalebars=2cm.

**Fig. S16.**
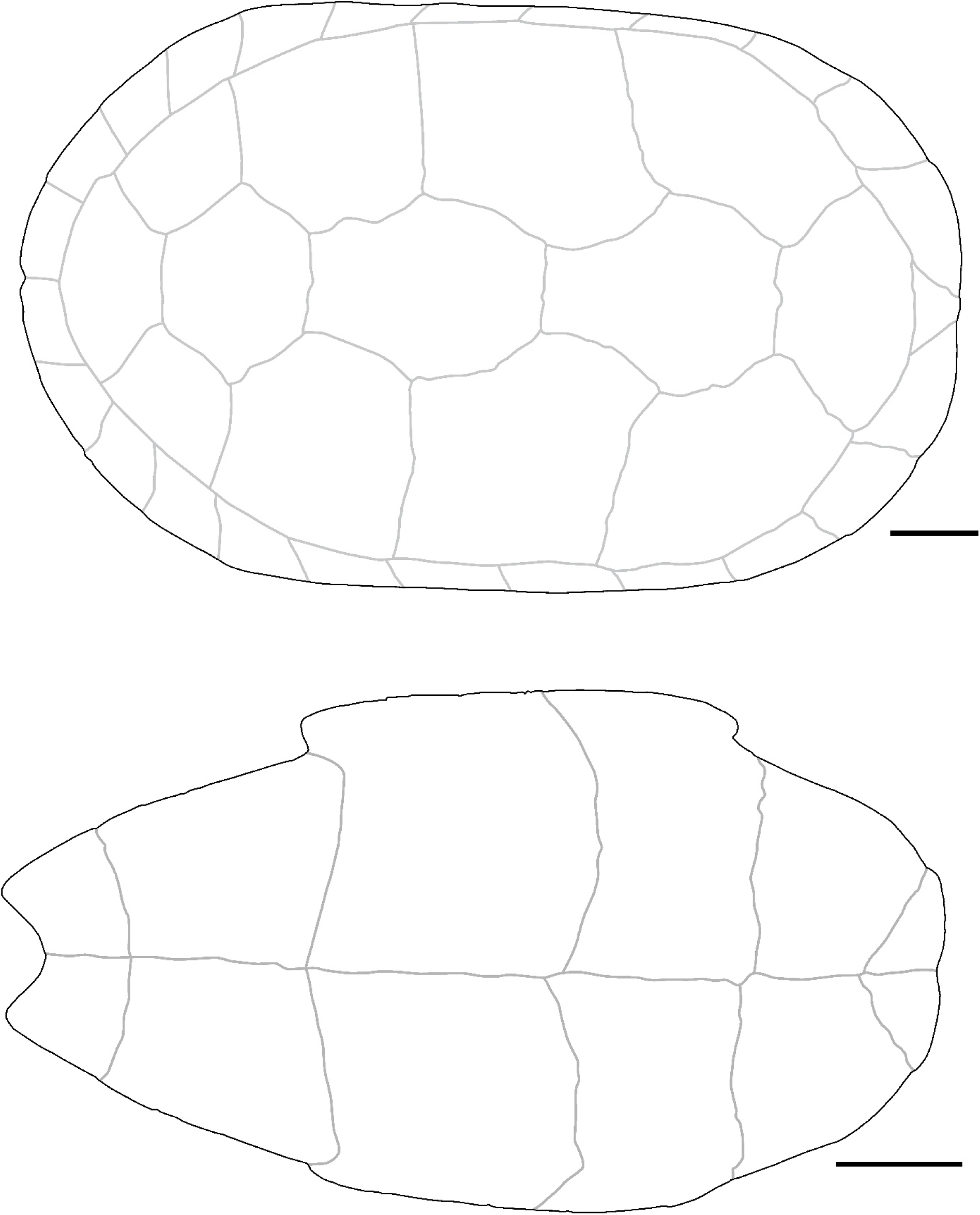
Carapace of an adult specimen of *Orlitia borneensis* Gray, 1873 (UF 152476) in dorsal (upper), and ventral (bottom) views. Scalebars=5cm.

**Fig. S17.**
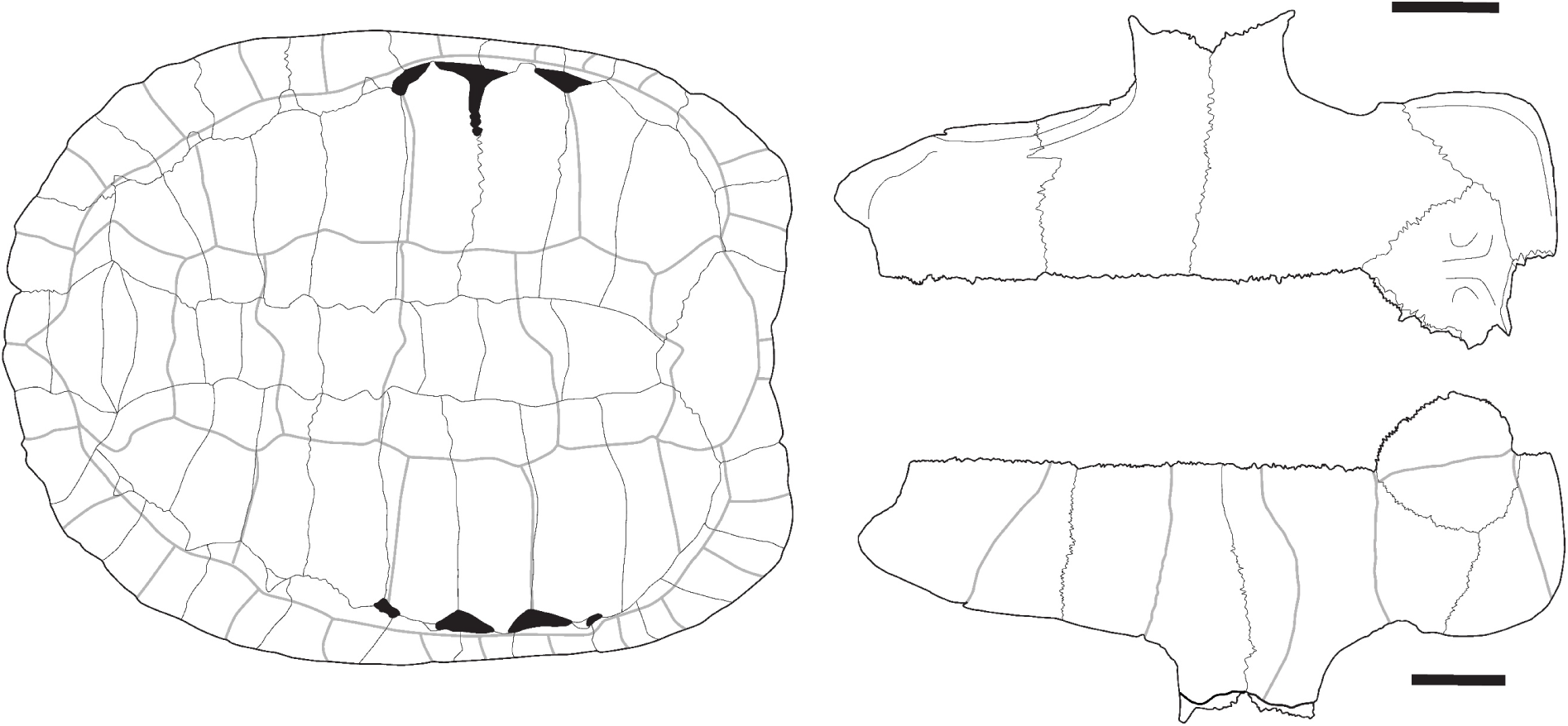
Carapace (UF 57111) and plastron (UF 56669) of *Platysternon megacephalum* Gray, 1831. Carapace in dorsal view (left) and plastron in dorsal (upper right) and ventral (bottom right) views. Scalebars=2cm.

**Fig. S18.**
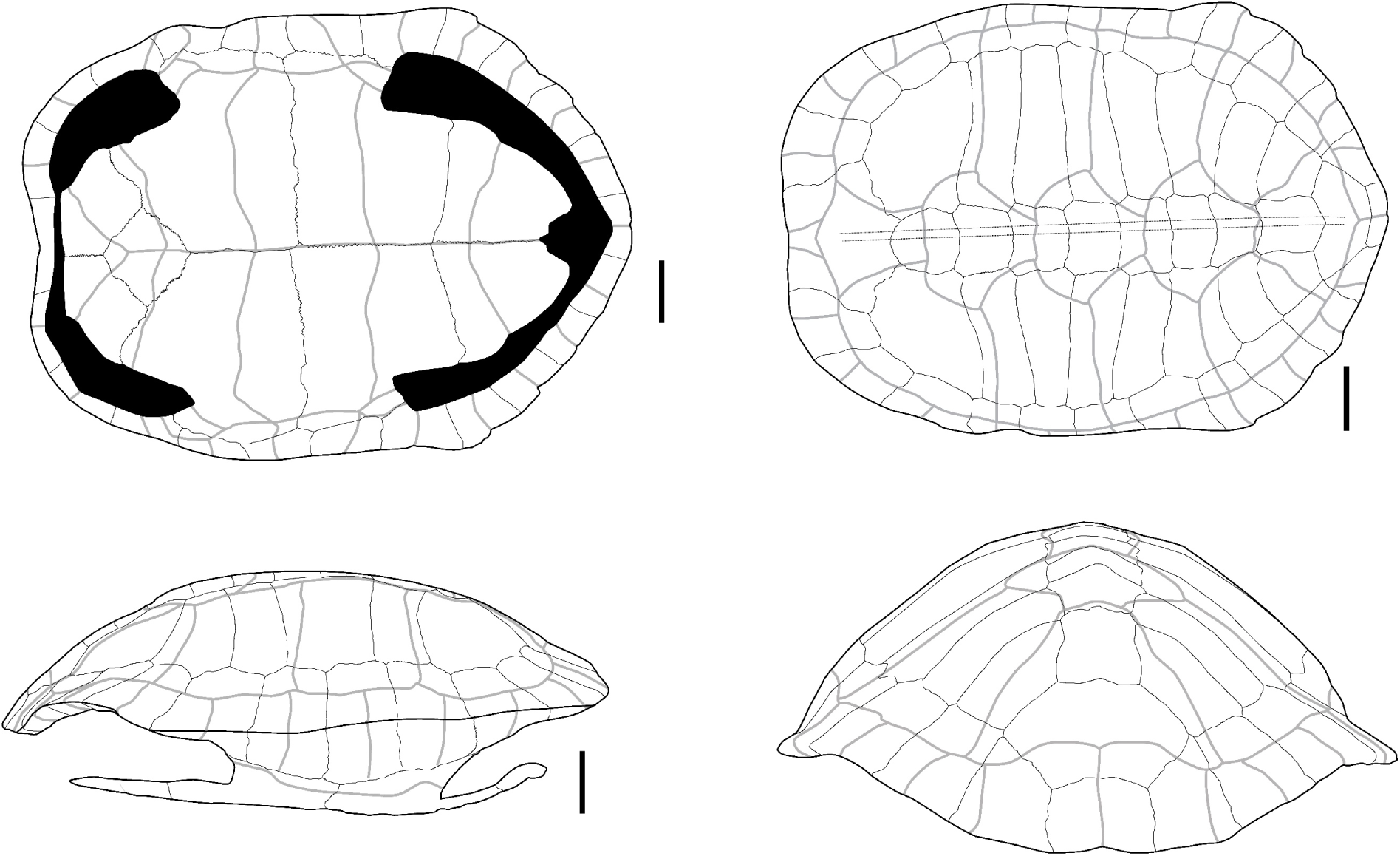
Carapace of *Siebenrockiella crassicollis* (Gray, 1830) (CUMZ-R-TG551) in ventral (upper left), lateral (bottom left), dorsal (upper right), and posterior (bottom right) views. Scalebars=2cm.

## Notes

### Competing Interest Statement

The authors have declared no competing interest.

### Summary of Updates

Addition of the link to the recommendation of PCI Paleontology.

